# Embryonic Vitamin D Deficiency Programs Hematopoietic Stem Cells to Induce Type 2 Diabetes

**DOI:** 10.1101/2022.09.08.507174

**Authors:** Jisu Oh, Amy E. Riek, Kevin T. Bauerle, Adriana Dusso, Kyle P. McNerney, Ruteja A. Barve, Isra Darwech, Jennifer Sprague, Clare Moynihan, Rong M Zhang, Ting Wang, Xiaoyun Xing, Daofeng Li, Richard D. Head, Monika Bambouskova, Marguerite Mrad, Alejandro Collins, Mark S. Sands, Carlos Bernal-Mizrachi

## Abstract

Environmental factors may alter the fetal genome to cause metabolic diseases. It is unknown whether embryonic immune cell programming impacts the risk of type 2 diabetes in later life. We demonstrate that transplantation of fetal hematopoietic stem cells (HSCs) made vitamin D deficient in utero induces diabetes in vitamin D-sufficient mice. Vitamin D deficiency epigenetically suppresses *Jarid2* expression and activates the *Mef2*/*PGC1a* pathway in HSCs, which persists in recipient bone marrow, resulting in adipose macrophage infiltration. These macrophages secrete miR106-5p, which promotes adipose insulin resistance by repressing PIK3 catalytic subunit alpha and AKT signaling. Vitamin D-deficient monocytes from human cord blood have comparable *Jarid2/Mef2/PGC1a* expression changes and secrete miR-106b-5p, causing adipocyte insulin resistance. These findings suggest that vitamin D deficiency during development has epigenetic consequences impacting the systemic metabolic milieu.

## Introduction

Nearly 100 million Americans have either type 2 diabetes mellitus (T2DM) or prediabetes as a manifestation of insulin resistance (IR) ^1^. Because rates of T2DM and its complications are increasing at such an alarming rate, America’s youth, for the first time in modern history, may have a shorter life expectancy than their parents ^2^. Although chronic inflammation plays a crucial role in the development of insulin resistance, the predisposing factors that alter immune cells are still unclear. Since the early 1980s, several historical cohort studies in diverse populations have provided evidence for the “developmental origins of adult disease hypothesis”, which posits that environmental factors in utero or during the early postnatal period program patterns of infant growth that result in increased susceptibility to IR and obesity later in life ^3, 4, 5^. Therefore, identifying such factors and the tissues affected by the epigenetic program initiated in utero are key to developing treatment strategies for patients and preventative therapies for future generations. During embryogenesis, the genome undergoes a critical period of reprogramming in response to environmental stimuli ^6^. While this process of genome-wide epigenetic reprogramming can provide an evolutionary benefit by facilitating rapid adaptation of the phenotype to the environment, it can also induce lifelong maladaptive changes which predispose individuals to IR and obesity later in life ^7, 8^. Early studies examining the regulation of metabolic disease by the epigenome from diverse exposures such as maternal under-or overnutrition were focused on metabolically active tissues such as the liver, endocrine pancreas, and adipose tissue. However, despite the fact that immune cells might be a unifying mechanism causing metabolic disease, there is a lack of studies that identify the epigenetic signature(s) of immune cells programmed during embryogenesis that predisposes to adult onset of IR and diabetes.

Changes to chromatin structure and function during embryogenesis are critical in determining appropriate gene expression patterns that drive distinct cell lineages ^9^. Epigenetic modifications such as DNA CpG methylation and covalent modifications of histones, can alter gene expression, often by structural modifications in the absence of changes in DNA sequence ^10^. Modulation of DNA methylases (DNMT) along with histone-modifying enzymes regulating chromatin function have been previously linked to IR ^11^. Deletion of H3K9-specific demethylase Jhdm2a induces obesity, hyperlipidemia, and IR in mice by repressing adipose PPARα and UCP-1 expression ^12^. Mice lacking DNMT3a are protected from diet-induced IR and glucose intolerance ^13^, and pharmacological inhibition of DNMTs relieves DNMT1 repression of adipose adiponectin expression, improving IR in obese mice ^14^. These data suggest that genetic and pharmacologic inhibition of chromatin-opening enzymes promotes global metabolic dysregulation and IR ^15^. Interestingly, studies in monozygotic twins indicated a direct correlation between global DNA methylation in peripheral blood leukocytes and the severity of IR, implicating the epigenetic modification of immune cells in the development of diabetes ^16^. Moreover, epidemiological studies have demonstrated an association between inactivating mutations of the epigenetic enzyme TET2, which induces clonal myeloid and lymphoid expansion, and the development of T2DM and cardiovascular disease in humans ^17, 18^. In mice, a causal relationship between immune cell dysfunction and metabolic disease has been illustrated by bone marrow transplantation from donor mice with inactivating mutations of TET2 to wild-type recipient mice, which was sufficient to induce macrophage adipose infiltration, pro-inflammatory cytokine IL-1β secretion, and obesity-related IR ^19^. Thus, discovering environmental factor(s) regulating the enzymes controlling the chromatin state in immune cells during embryogenesis could be critical to the prevention of chronic inflammatory diseases such as diabetes.

In the U.S., 80% of pregnant African-American females and 60% of pregnant Caucasian females are vitamin D-deficient or insufficient ^20^. Vitamin D deficiency [VD(−)] is associated with low birth weight and small for gestational age, with increased susceptibility to obesity, IR, and diabetes later in life ^21, 22, 23^. Murine studies confirm that in utero vitamin D deficiency results in offspring systemic inflammation, hepatic steatosis, excess adiposity, and IR that persist despite vitamin D supplementation after birth, implying that vitamin D deficiency during gestation induces epigenetic programming ^24, 25, 26, 27, 28^. However, the tissue(s) carrying the underlying cellular program to cause offspring IR has remained elusive.

Multiple studies strongly suggest the role of vitamin D receptor (VDR) in hematopoiesis. VDR knockout mice have persistent changes in lymphocytic and myelocytic function and cytokine profiles, suggesting the importance of VDR signaling for immune cell programming during embryogenesis ^29^. Vitamin D is already known to regulate multiple components of the epigenetic machinery, though the net effects of this regulation are mixed depending upon the system studied ^30^. Polycomb group (PcG) proteins, made up of the initiation complex PRC2 and maintenance complex PRC1, play an important role in the maintenance of transcriptional repression of target genes through chromatin modifications ^31^. PRC2 requires Jarid2 for its repressive activity, and both affect proliferative and self-renewal capacities of hematopoietic stem cells to influence their immune program ^32^. Prior studies have linked active vitamin D to upregulation of murine macrophage *Jarid2* expression, but the effects of this interaction on the immune cell epigenome as it contributes to metabolic disease are unknown ^30^.

Our previous studies indicated that macrophage-specific VDR deletion is sufficient to induce IR and hypertension by promoting a pro-inflammatory macrophage phenotype in metabolic tissues, suggesting that altered VDR signaling in immune cells during embryogenesis programs proinflammatory immune cells that to cause IR in the offspring ^28, 33^. In this study, by using transplantation of fetal hematopoietic stem cells (HSCs) exposed to VD deficiency in utero into VD-sufficient mice, we identified an epigenetic program of a single, non-metabolically active tissue compartment that is mitotically stable and sufficient to induce type 2 diabetes.

## Results

### Hematopoietic Stem Cells from Vitamin D Deficient Embryos Transplant Insulin Resistance

To determine whether the fetal immune cell program induced by VD deficiency is sufficient to cause IR in different mouse backgrounds, C57BL/6 and C57BL/6-LDLR^−/−^GFP^+/−^ (a model of diet-induced IR) female mice were fed a vitamin D-deficient or sufficient diet four weeks before pregnancy. We confirmed that the dams were vitamin D deficient at mid-gestation [25(OH)D 9±3 vs. 35±2 ng/mL]. There were no differences in dam or fetus weights, food intake, serum calcium, glucose, or lipids between VD(−) and VD(+) dams **(Supplementary Fig. 1A-E)**. We isolated fetal (E13) liver HSCs from embryos obtained from VD(−) and VD(+) dams in the C57BL/6 and C57BL/6-LDLR^−/−^GFP^+/−^ background. HSCs were then transplanted into genotype-matched eight-week-old VD-sufficient mice. Eight weeks after transplantation, both groups of primary recipients had similar weights, and 90% of peripheral blood cells were of donor origin **(Supplementary Fig. 2A-D)**. Mice of both sexes and genetic backgrounds receiving VD(−) HSCs demonstrated fasting hyperglycemia, impaired glucose tolerance as determined by intraperitoneal glucose tolerance testing (GTT), and IR as determined by insulin tolerance testing (ITT) **(Fig. 1A and B; Supplementary Fig. 3A and B)**. To determine if the IR phenotype was transmitted by pluripotent HSCs, bone marrow from the primary recipients was transplanted into VD(+) secondary recipients. At six months post-transplant, both primary and secondary recipients maintained a stable IR phenotype **(Fig. 1C-F)**. LDLR deletion did not affect the IR phenotype induced by VD(−) HSCs in either primary or secondary recipients (**Supplementary Fig. 3A-D)**. Transfer of VD(−) HSCs did not alter recipient insulin levels after glucose challenge (**Supplementary Figure 4A-C)**. These data confirm that in utero VD deficiency induces an HSC program that confers persistent IR in VD sufficient primary and secondary transplant recipients.

**Figure 1.**
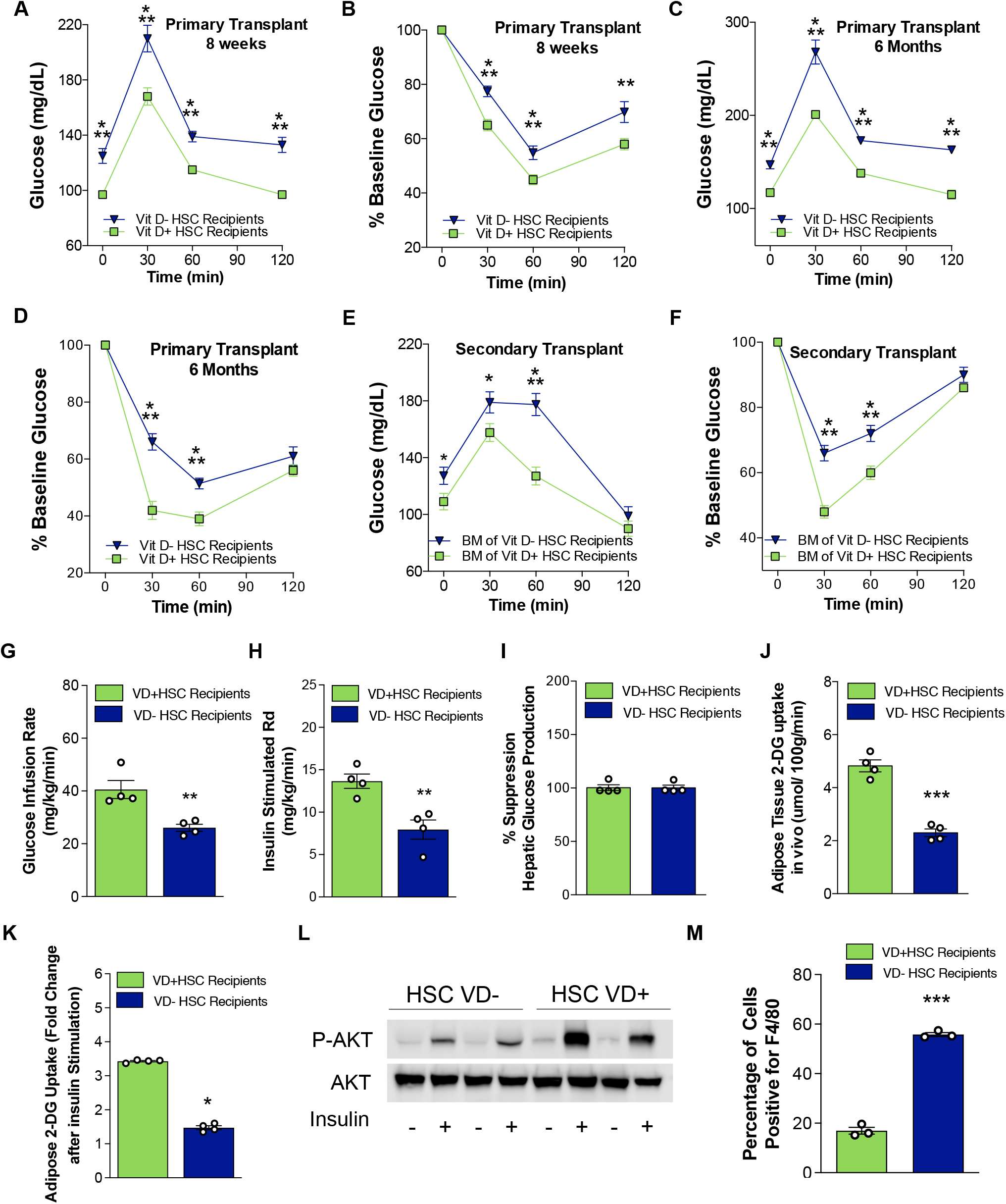
In utero VD deficiency reprograms HSCs to transfer IR. **(A-J)** Vitamin D sufficient CD45.2^+^ C57BL6 mice were transplanted with VD(−) or VD(+) FL-HSCs from CD45.1^+^ C57BL6 mice (primary; n=20/group), then these primary recipients were used as BM transplant donors for vitamin D sufficient mice (secondary; 20/group). Glucose and insulin tolerance tests were performed at **(A and B)** 8 weeks (n=18-20/group) and **(C-D)** 6 months post-primary-transplant (n=15-16/group) and **(E and F)** 8 weeks post-secondary-transplant (n=12/group). Hyperglycemic-euglycemic clamps were conducted in primary transplant recipients after 8 weeks (n=4/group). Data are reported as **(G)** glucose infusion rate, **(H)** insulin-stimulated glucose disposal rate (Rd), **(I)** change in hepatic glucose production, and **(J)** insulin-stimulated 2-DG uptake in adipose tissue. **(K-M)** Periogonadal white adipose tissue was isolated from VD(−) or VD(+) FL-HSC recipients. **(K)** *Ex vivo* insulin-stimulated 2-DG uptake (n=4/group). **(L)** Western blot analysis of phospho-and total AKT levels following insulin stimulation. **(M)** Percentage of F4/80-positive cells by manual counting of 15 light microscopy fields under 20x objective per mouse (n=3/group). Data presented as mean ± SEM. *p<0.05; **p<0.01; ***p<0.001 vs. VD(+) FL-HSCs recipients by two-tailed unpaired t-test.

Hyperglycemic-euglycemic clamping performed eight weeks after the primary transplant showed that peripheral IR was induced in C57BL6 VD(−) HSC recipients under conditions of vitamin D sufficiency. (**Fig. 1G-I**). Insulin-stimulated uptake of 2-deoxyglucose (2-DG) at the end of the clamp identified perigonadal fat rather than the muscle as the primary insulin-resistant tissue (**Fig. 1J, Supplementary Fig. 4D and E**). These findings in perigonadal adipose tissue of VD(−) HSC recipients were supported *ex vivo* by a reduction in insulin-stimulated 2-DG uptake and phospho-AKT, as well as increased macrophage infiltration (**Fig. 1K-M**). Together, these results suggest that in utero VD deficiency induces an HSC program that promotes adipose macrophage infiltration to cause IR.

### Fetal Vitamin D Deficiency Represses Jarid2 Expression

To elucidate the HSC program associated with VD deficiency, we performed a multi-omic analysis of mRNA and miRNA expression. Transcriptome analysis showed consistent upregulation of 391 genes and downregulation of 657 genes in the BM of VD(−) vs. VD(+) HSC transplant recipients at 8 weeks post-transplant (GEO:GSE158763) (**Fig. 2A)**. Enrichment pathway analysis identified the Jarid2 pathway signature as the most significantly activated (**Fig. 2B, Supplementary Table 1**). Jarid2, a histone methyltransferase that is part of the polycomb repressive complex 2 (PRC) and critical for immune cell differentiation ^31^, was downregulated, resulting in the expected activation of downstream genes involved in metabolic function, specifically myocyte enhancer factor 2 (Mef2) and its coactivator, peroxisome proliferator-activated receptor gamma coactivator 1-alpha (PGC-1*α*) (**Fig. 2C-F; Supplementary Fig. 5A)** ^34, 35, 36^. Indeed, silencing of *Jarid2* expression in peritoneal macrophages also activated the *Mef2/PGC1α* pathway (**Fig. 3A, Supplementary Fig. 5B**). Alterations in the *Jarid2/Mef2/PGC1α* pathway were also present in donor VD(−) HSCs and recipient macrophages despite normal plasma VD levels in recipient mice **(Fig. 3B and C, Supplementary Fig. 5C and D)**, suggesting that this genetic program activated in VD (-) HSCs was persistent in immune cells from VD(−) HSC-transplanted mice. These findings are consistent with previous evidence that VD stimulates *Jarid2* expression ^30^. To determine if this immune cell program was transmitted through changes in the immune cell epigenome, pups exposed to VD deficiency in utero were provided postnatal VD supplementation. Indeed, *Jarid2* expression was not restored with VD supplementation, indicating that epigenetic modifications resulting from in utero VD deficiency are responsible for the observed changes in macrophage *Jarid2* expression that contribute to *Mef2/PGC1α* pathway activation (**Fig. 3D**). We hypothesized that the gene expression changes leading to stable reprogramming of the *Jarid2/Mef2/PGC1α* pathway were a consequence of changes in *Jarid2* methylation status. Therefore, we performed targeted next-generation sequencing (NGS) methylation assays to interrogate the DNA methylation status of 69 CpG sites in the 5’ upstream through 3’ UTR regions of the mouse *Jarid2* gene in the BM from VD(−) and VD(+) HSC recipients (**Supplementary Table 2**). We found an increase in the methylation status of several CpG sites within or near putative *Jarid2* enhancers in the BM from VD(−) HSC recipients **(Fig. 3E)**.

**Figure 2.**
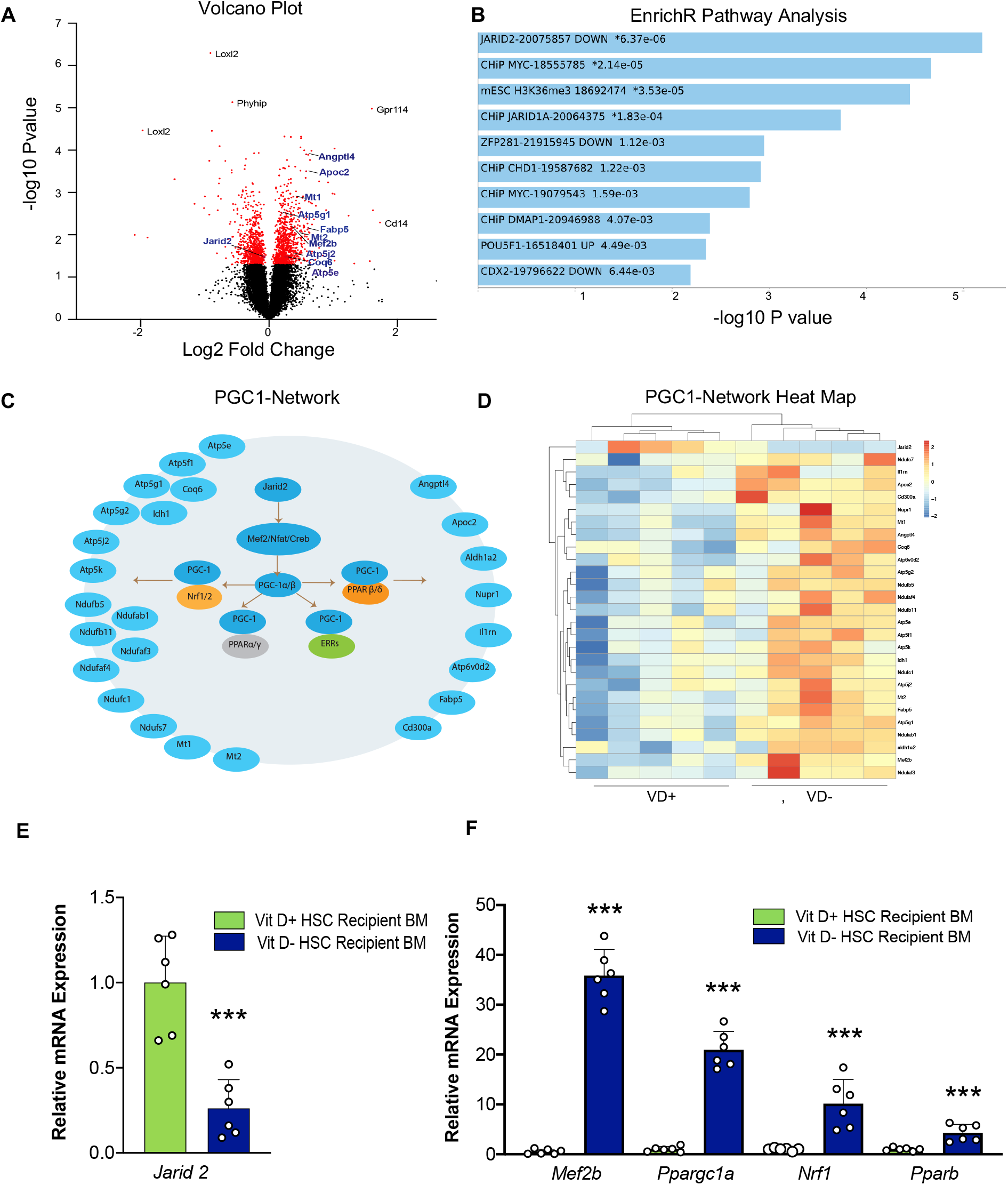
Top genes, networks, and pathways identified in transcriptome analysis of bone marrow from recipient mice transplanted with HSCs isolated from embryos from VD sufficient and deficient dams. VD(−) vs. VD(+) FL-HSCs were transplanted into VD(+) mice, and global mRNA expression was evaluated by microarray in recipient BM cells at 16 weeks post-transplant. (A) Volcano plot showing top differentially expressed genes. Red dots indicate those array probes with p<0.05. Black represents the non-significant probes. Genes of the Jarid2-PGC1-MEF2 pathway are highlighted in blue. (B) Genes with significant changes were used for manual and automated pathway analysis. The figure shows EnrichR (PMID 23586463; 27141961; 33780170) top pathway hits from ESCAPE database. Asterisks indicate those pathways that pass multiple testing corrections. (C) Illustration of the Jarid2-MEF2-PGC1 network and the target genes that are differentially expressed in the array data. (D) Heat map table showing normalized gene expression for Jarid2, MEF2-PGC1 target genes. Red indicates upregulation, and blue indicates downregulation. (E and F) Quantitative RT-PCR in FL-HSC transplant recipient BM to confirm expression changes in Jarid2 and PGC1α network-related genes.

**Figure 3.**
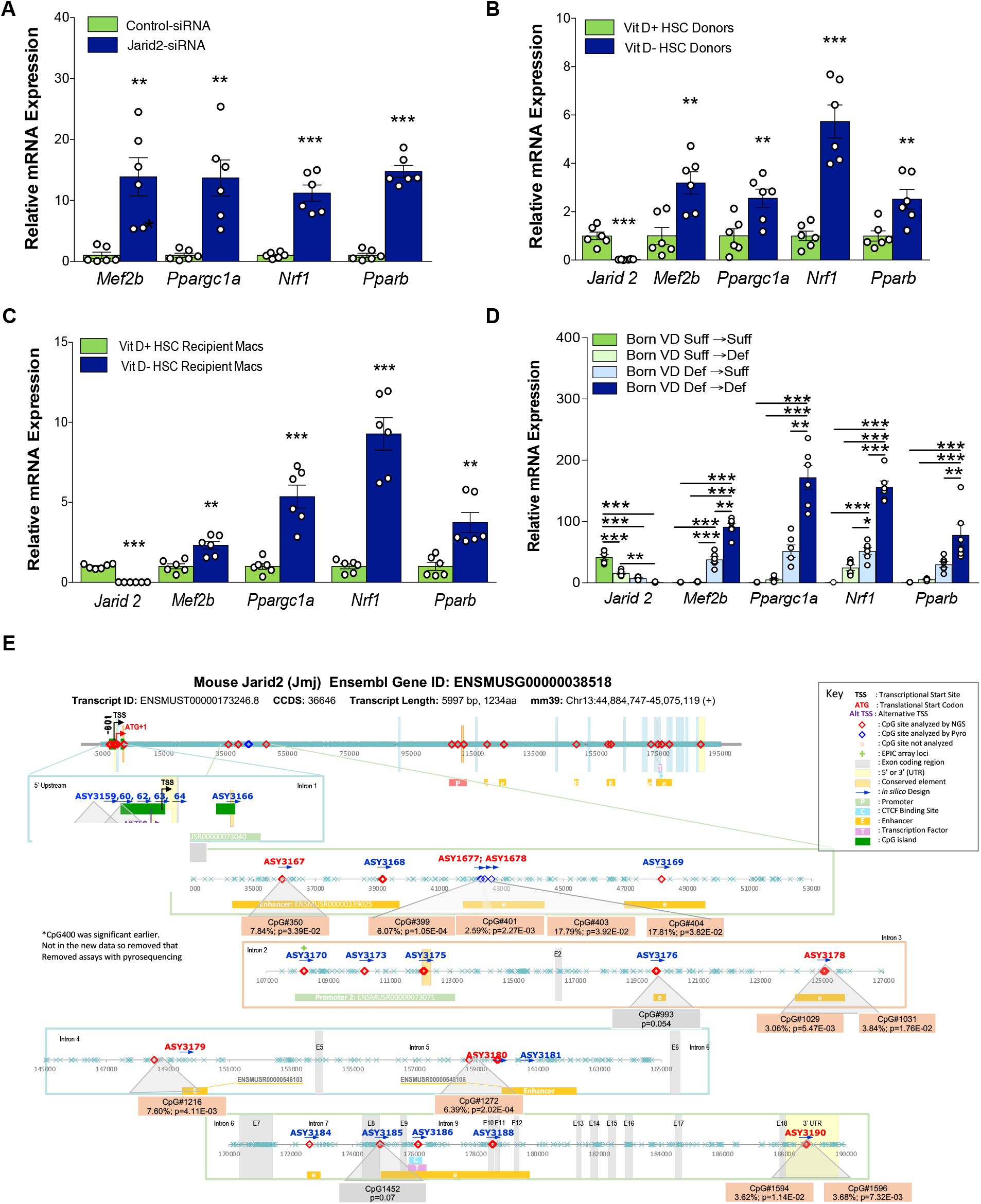
In utero VD deficiency epigenetically suppresses Jarid2 expression in macrophages, resulting in persistent *PGC1α* network upregulation. **(A)** Quantitative RT-PCR in peritoneal macrophages (n=6/group) transfected with Jarid2-siRNA vs. control-siRNA. **(B and C)** Quantitative RT-PCR in donor FL-HSCs and peritoneal macrophages from FL-HSC transplant recipients (n=6/group). **(D)** Quantitative RT-PCR in macrophages to analyze expression of Jarid2 and PGC1α network-related genes in mice born VD(−) or VD(+) then fed VD(−) or VD(+) diet post-natally (n=6/group). **(E)** Targeted next-generation sequencing methylation assays were performed to interrogate the 69 CpG sites in the 5’ Upstream through 3’ UTR regions of the mouse *Jarid2* gene, including 2 promoter regions and several enhancers, in BM isolated from VD(−) and VD(+) FL-HSCs recipients at 16 wks (n=4/group). Shown is the gene structure of mouse *Jarid2* and the regions throughout the gene where the CpG methylation was interrogated. The methylation assays highlighted in red demonstrate a significant difference (p <0.05) between the two cohorts. The CpGs in these regions are shown in orange boxes. The % methylation increase and P values are reported. The grey boxes are those CpGs that have suggestive changes.

### Jarid2/PCG1α Programming Induces Insulin Resistance

To test whether the macrophage epigenetic program activated in VD(−) HSCs induces adipose IR, we silenced *Jarid2* expression in peritoneal macrophages from VD (+) HSC recipients. Silencing of macrophage *Jarid2* augmented inflammatory cytokine secretion (TNF*α*, IL-1β, IL-6) and adipose IR in co-culture experiments with 3T3-L1 adipocytes (**Fig. 4A and B**). Conversely, PGC1*α* deletion in peritoneal macrophages from VD(−) HSC recipients improved adipose IR in co-culture experiments with 3T3-L1 adipocytes and suppressed inflammatory cytokine release (**Fig. 4C and D**), suggesting that activation of this inflammatory pathway in macrophages by in utero VD deficiency may be responsible for adipose IR.

**Figure 4.**
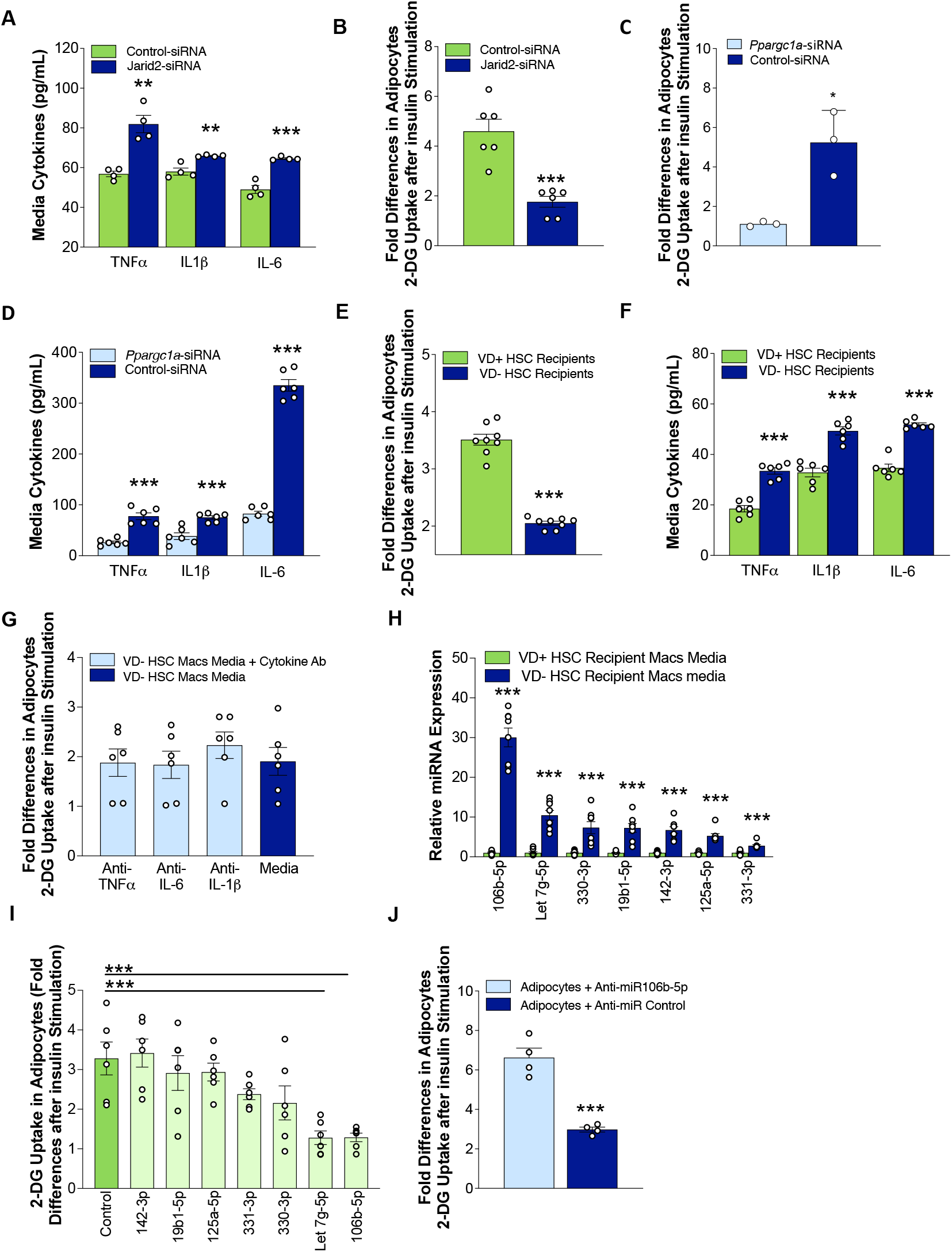
Activation of the Jarid2/Mef2/PGC1*α* immune cell program by in utero VD deficiency promotes proinflammatory cytokine and miRNA release. **(A**) Secreted cytokine levels in media (n=4/group) in Jarid2-siRNA vs. control-siRNA-transfected peritoneal macrophages from VD(+) FL-HSCs recipients. **(B)** Insulin-stimulated 2-DG uptake in 3T3-L1 adipocytes co-cultured with peritoneal macrophages from VD(+) FL-HSCs recipients transfected with Jarid2-siRNA vs. control-siRNA (n=6/group). **(C)** Insulin-stimulated 2-DG uptake in 3T3-L1 adipocytes co-cultured with peritoneal macrophages from VD(−) FL-HSCs recipients transfected with *Ppargc1a*-siRNA vs. control-siRNA. **(D)** Secreted cytokine levels in media (n=6/group) from *Ppargc1a*-siRNA vs. control-siRNA-transfected peritoneal macrophages from VD(−) FL-HSCs recipients. **(E)** Insulin-stimulated 2-DG uptake in 3T3-L1 adipocytes co-cultured with SVF macrophages from VD(−) or VD(+) HSC recipients (n=8/group). **(F)** Secreted cytokine levels in media (n=6/group) from SVF macrophages from VD(−) or VD(+) HSC recipients. **(G)** Insulin-stimulated 2-DG uptake in 3T3-L1 adipocytes co-cultured with SVF macrophages from VD(−) HSC recipients treated with or without cytokine-neutralizing antibodies (n=6/group). **(H)** Relative miRNA content in exosomes secreted by peritoneal macrophages from VD(−) vs. VD(+) FL-HSCs recipients (n=8/group). **(I)** Insulin-stimulated 2-DG uptake in 3T3-L1 adipocytes after transfection with miR mimics (n=6/group). **(J)** Insulin-stimulated 2-DG uptake in 3T3-L1 adipocytes transfected with miR-106b antagomir or control and cultured in conditioned macrophage media from VD(−) HSC recipients (n=4/group). Data presented as mean ± SEM. **p<0.01; ***p<0.001 by two-tailed unpaired t test. **(I)** *p<0.05; **p<0.01; ***p<0.001 by one-way ANOVA followed by Tukey’s multiple comparison test.

To investigate how adipose tissue macrophages (ATM) isolated from VD(−) and VD(+) HSC recipients induce IR in adipose tissue, we co-cultured 3T3-L1 adipocytes in transwell chambers with stromal vascular fraction macrophages. Adipocytes co-cultured with VD(−) HSC recipient macrophages had lower insulin-stimulated 2-DG uptake compared to those cultured with macrophages from VD(+) HSC recipients (**Fig. 4E**). Macrophages from VD(−) HSC recipients demonstrated increased secretion of TNF*α*, IL-1β, and IL-6 (**Fig. 4F**), but the addition of cytokine-neutralizing antibodies did not improve the adipocyte IR in co-culture experiments with 3T3-L1 adipocytes (**Fig. 4G**), suggesting that the IR phenotype may not be cytokine-driven.

### Macrophage miR106b-5p Secretion Causes Adipocyte Insulin Resistance

Recently, microRNAs (miRNAs) have shown to regulate chronic inflammation and IR ^37, 38^. Numerous immature miRNAs were downregulated within BM cells from VD(−) HSC recipients (**Supplementary Table 3**). However, the corresponding VD(−) HSC recipient peritoneal macrophage media had increased levels of mature miRNAs, suggesting enhanced macrophage miRNA maturation and secretion. Similar to our previous finding in mice with macrophage-specific VDR deletion ^33^, the most highly secreted miRNA from VD(−) HSC recipient peritoneal macrophage was miR106b-5p (**Fig. 4H**). Transfection of mouse 3T3-L1 adipocytes with miRNA mimics of the most abundant secreted macrophage miRNAs identified in VD (-) recipient macrophages showed that miR-106b-5p induced the most significant adipocyte IR (**Fig. 4I)**. Moreover, adipocytes transfected with miR-106b-5p antagomir and exposed to conditioned media from peritoneal macrophages from VD(−) HSC recipients demonstrated improved insulin sensitivity (**Fig. 4J)**.

To clarify how miR-106b-5p causes adipose IR, we transfected 3T3-L1 adipocytes with a miR-106b-5p mimic to determine if miR-106b-5p regulates insulin-stimulated PIK3 signaling activation. Indeed, miR-106b-5p mimic transfection reduced transcript levels of PIK3CA, the primary insulin-responsive PI3K isoform in adipocytes ^39^, and the 3-phosphoinositide dependent protein kinase 1 (PDPK1), which is responsible for AKT activation. Of note, transcript levels of PIK3CB were not altered by miR-106b-5p mimic transfection. Western blot analysis confirmed decreased protein levels of PIK3CA and PDPK1 (**Fig. 5A-C)**. In contrast, transfection of 3T3-L1 adipocytes with miR-106b-5p antagomir prevents suppression of PIK3CA and PDPK1 expression induced by macrophage media from VD(−) HSC recipient **(Fig. 5D-F)**. These results suggest that miR106b-5p induced-downregulation of PIK3CA/PDK1/AKT signaling pathway is a primary mechanism driving adipose IR.

**Figure 5.**
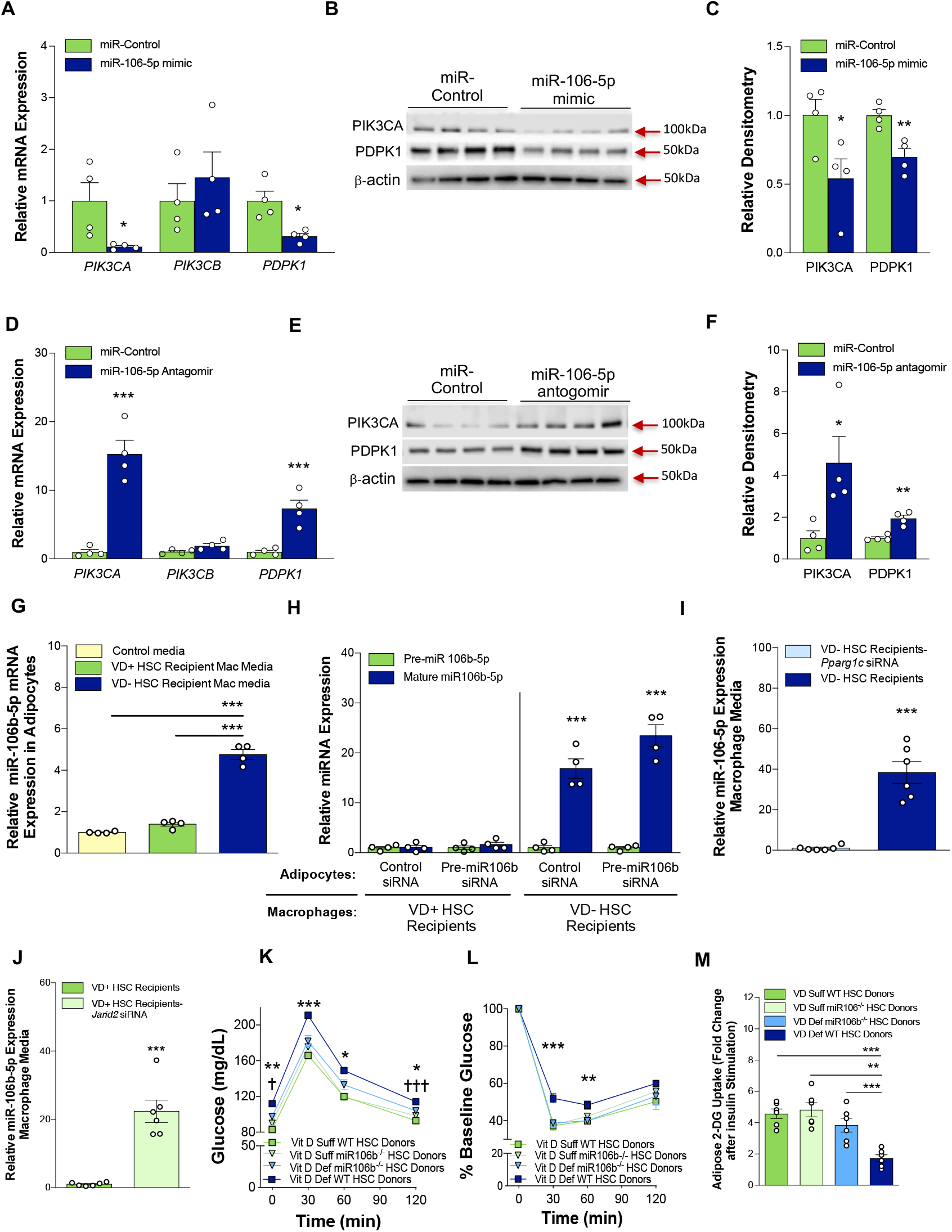
Macrophage miR-106b-5p mediates in utero VD deficiency-induced adipocyte IR. **(A-C)** Quantitative RT-PCR, western blot analysis, and densitometry (normalized to β-actin protein levels) of insulin signaling pathway in 3T3L1 cells after transfection with miR106b-5p mimic vs. control mimic. **(D-F)** Quantitave RT-PCR, western blot analysis and densitometry (normalized to β-actin protein levels) of insulin signaling pathway in 3T3L1 cells cultured in conditioned media from VD(−) HSC recipient macrophages after transfection with anti-miR-106b or control (n=4/group). **(G)** miR-106b-5p expression in adipocytes cultured in conditioned media from macrophages isolated from VD(−) or VD(+) HSC recipients (n=4/group). **(H)** Pre-and mature-miR-106b-5p abundance in 3T3-L1 adipocytes transfected with pre-miR-106b-siRNA vs. control-siRNA then cultured in conditioned media from macrophages isolated from VD (-) or VD(+) HSC recipients (n=4/group). Peritoneal macrophage media miR-106b-5p expression from **(I)** VD(−) HSC recipient macrophages with or without *Ppargc1a*-siRNA, and **(J)** VD(+) HSC recipient macrophages with or without Jarid2-siRNA (n=6/group). **(K-M)** Fetal HSCs from WT or miR-106b^−/−^ animals under VD(−) or VD(+) conditions were transplanted into VD(+) WT recipients. **(K)** Glucose tolerance tests and **(L)** insulin tolerance tests (n=8/group). **(M)** Insulin-stimulated 2-DG uptake in 3T3-L1 adipocytes co-cultured with peritoneal macrophages from WT or miR-106b^−/−^ animals transplanted with VD(−) or VD(+) HSCs (n=6/group). Data presented as mean ± SEM. **(A**,**C**,**D**,**F**,**H**,**I**,**J)** *p<0.05; ***p<0.001 by two-tailed unpaired t test. **(G and M)** *p<0.05; **p<0.01; ***p<0.001 by one-way ANOVA followed by Tukey’s multiple comparison test. **(K and L)** *p<0.05; **p<0.01; ***p<0.001 VD-WT vs. all and ^†^p<0.05; ^†††^p<0.001 for VD+ WT vs. VD-miR-106b^−/−^ by one-way ANOVA followed by Tukey’s multiple comparison test.

Adipocytes expressed little miR-106b-5p at baseline. However, adipocyte exposure to conditioned macrophage media from VD(−) HSC recipients increased mature miR-106b-5p abundance by 7-fold compared to VD(+) HSC recipient conditioned macrophage media **(Fig. 5G)**. To define the cell responsible for producing mature miR-106b-5p, adipocytes were co-cultured with macrophages from VD(−) and VD(+) HSC recipients and transfected with a pre-miR-106b siRNA to inhibit endogenous adipocyte miR-106b production. Increased abundance of mature miR-106b-5p in the media and exacerbation of IR in adipocytes was noted despite pre-miR-106b siRNA transfection when adipocytes were exposed to media from VD(−) HSC recipients, confirming that macrophages are the source of the miR-106b-5p that drives the IR phenotype **(Fig. 5H, Supplementary Fig. 6)**. Finally, we found that silencing *Ppargc1a* in VD(−) HSC recipient macrophages suppressed miR-106b-5p secretion (**Fig. 5I**), while silencing *Jarid2* in VD(+) HSC recipient macrophages promoted miR-106b-5p secretion (**Fig. 5J**). Overall, these results indicate that the Jarid2/Mef2/PGC1*α* immune cell program induced by VD deficiency in utero regulates macrophage miR-106b-5p secretion.

**Figure 6.**
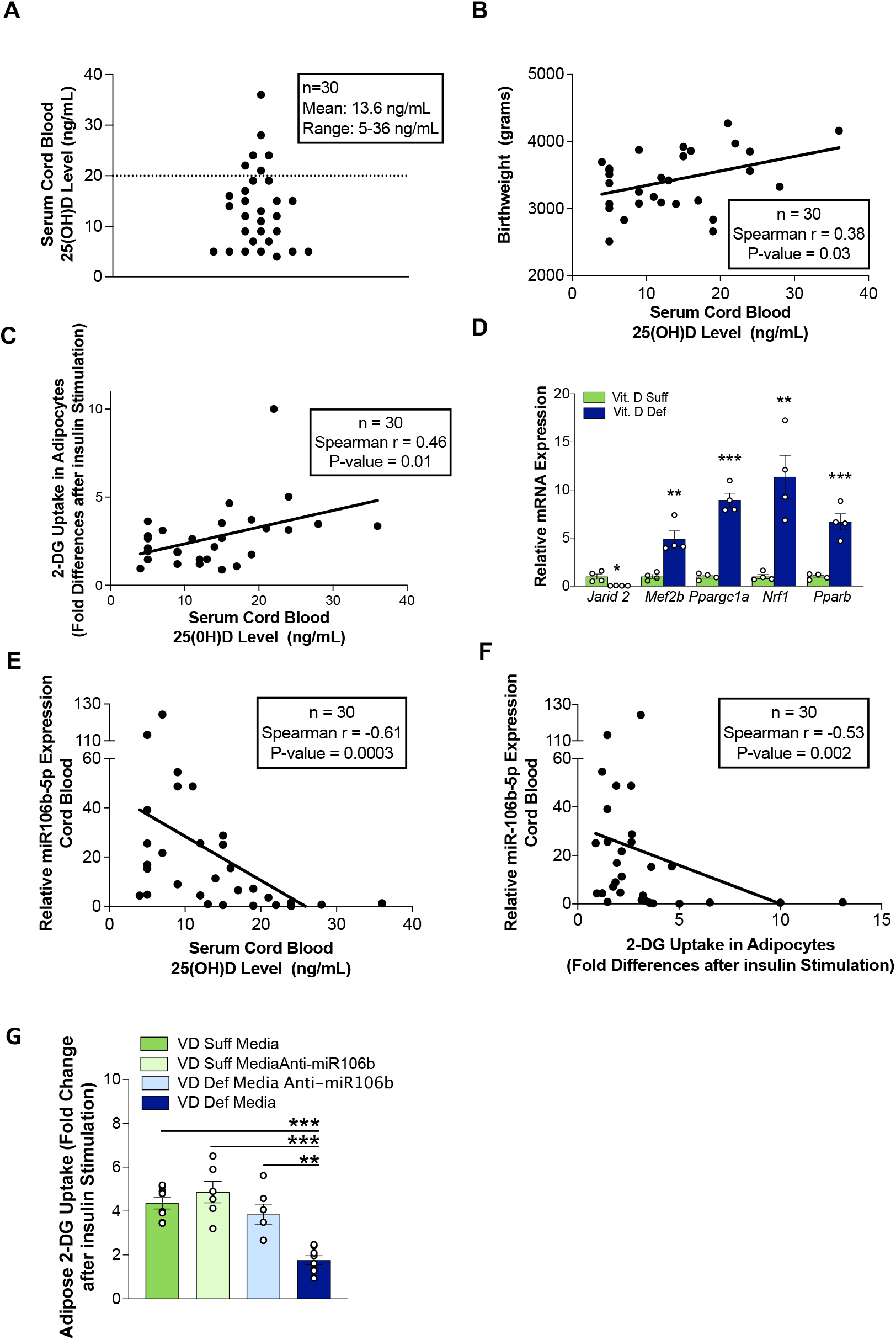
VD deficient cord blood monocytes induce adipocyte IR. **(A)** Cord blood serum 25(OH)D levels from 30 healthy pregnant women at delivery. **(B)** Correlation between cord blood serum 25(OH)D levels and birthweight. **(C)** Correlation between cord blood serum 25(OH)D levels and change in insulin stimulated 2-DG uptake in 3T3-L1 adipocytes cultured in conditioned media of cord blood monocytes. **(D)** Quantitative RT-PCR of mRNA expression of Jarid2/Mef2/PGC1α network-related genes from cord blood monocytes stratified by 25(OH)D level (25(OH)D <20 ng/mL and β20 ng/mL) (n=4/group). **(E)** Correlation between cord blood serum 25(OH)D level and serum miR-106b-5p expression (n=30). **(F)** Correlation between serum miR-106b-5p expression and insulin-stimulated 2-DG uptake in 3T3-L1 adipocytes cultured in conditioned media from cord blood monocytes (n=30). **(G)** Insulin-stimulated 2-DG uptake in 3T3-L1 adipocytes transfected with miR-106b-5p antagomir and cultured in conditioned media from blood monocytes with 25(OH)D <20 ng/mL or β20 ng/mL (n=6/group). Data presented as mean ± SEM. **(D)** *p<0.05; **p<0.01; ***p<0.001 by two-tailed unpaired t test. **(G)** **p<0.01; ***p<0.001 by one-way ANOVA followed by Tukey’s multiple comparison test.

To confirm the role of macrophage miR-106b-5p in the IR phenotype, we transplanted VD(−) or VD(+) HSCs from miR-106^−/−^ or miR-106^+/+^ mice into VD sufficient control mice. Recipients of VD(−) miR-106^−/−^ HSCs had improved glucose tolerance by GTT and improved adipose insulin sensitivity by ITT compared to VD(−) miR106^+/+^ HSC recipients (**Fig. 5K and L**). These findings were supported by co-culture experiments with 3T3-L1 cells and macrophages from VD(−) or VD(+), miR-106^−/−^ or miR-106^+/+^ HSCs recipients, where the absence of miR-106b-5p improved 3T3-L1 IR when cultured with macrophages from VD(−) HSC recipients (**Fig. 5M**), indicating that macrophage secretion of miR106b-5p is critical for the adipose IR phenotype induced by in utero VD deficiency.

### Cord Blood Monocytes from Vitamin D Deficient Subjects causes Adipocyte IR

To determine whether VD deficiency during pregnancy in humans induces similar HSC cell reprogramming, we analyzed 30 healthy pregnant women prior to delivery and their full-term infants (**Supplementary Table 4**). Two-thirds of newborns were VD-deficient [25(OH)D ≤20 ng/mL], and cord blood 25(OH)D levels directly correlated with newborn low birth weight (**Fig. 6 A and B**), consistent with previous studies ^40^. We found a direct correlation between cord blood 25(OH)D levels and 3T3-L1 adipocyte insulin sensitivity after exposure to conditioned media from cord blood monocyte (**Fig. 6C**). Additionally, cord blood monocytes from VD-deficient mothers had lower *Jarid2* expression and increased *Mef2/PGC1α* expression (**Fig. 6D**). Cord blood 25(OH)D levels also correlated inversely with cord blood plasma miR-106b-5p levels, which, as expected, was predictive of adipocyte IR **(Fig. 6E and F)**. Moreover, transfection of 3T3-L1 adipocytes with miR-106b-5p antagomir improved the insulin resistance induced by conditioned media from VD-deficient monocytes (**Fig. 6G**). These findings highlight the persistent effects of maternal vitamin D deficiency on offspring macrophage function to cause insulin resistance.

## Discussion

Despite recognition of the importance of environmental conditions in utero as contributing to adult disease, there is little data identifying which conditions during embryogenesis and which target tissues carry the epigenetic program that increases the susceptibility to IR in the offspring later in life. Vitamin D deficiency and insufficiency are highly prevalent at the time of delivery and concordant in both mothers and their neonates ^40^. This study provides direct evidence that vitamin D deficiency in utero is sufficient to induce stable epigenetic programming in HSCs that can be transplanted to induce IR in vitamin D-sufficient recipient mice and is not reversed with postnatal vitamin D supplementation. To our knowledge, this is the demonstration that an epigenetic modification of a single, non-metabolically active tissue compartment is sufficient to induce type 2 diabetes. VD deficiency epigenetically suppresses *Jarid2* expression and activates the *Mef2*/*PGC1a* pathway in fetal HSCs. This program persists in recipient monocytes/macrophages, promoting adipose macrophage infiltration and miR-106b-5p secretion to suppress adipose PIK3CA and AKT activity, causing IR. Lack of miR106b-5p prevents the VD(−) HSCs from adoptively transferring IR to recipients. These data strongly suggest that IR is caused by epigenetic reprogramming of immune cells induced by vitamin D deficiency in utero, leading to activation of the *Jarid2/Mef2/PGC1α*/*miR106b-5p* pathway both in human and mice.

Epigenetic memory is defined as the stable propagation of gene expression induced by an environmental or developmental stimulus and may be categorized as cellular, transgenerational, or transcriptional ^41^. Cellular memory refers to mitotically heritable transcriptional states and is elegantly illustrated in our model, in which an epigenetic program is induced by in utero VD deficiency in HSCs and then transmitted to the bone marrow and monocyte/macrophage lineages. Furthermore, the stability of this memory is highlighted by the secondary BM transplant experiments in which BM from VD deficient HSC recipients was capable of inducing IR in VD sufficient mice. Cellular memory requires Trithorax and PcG group proteins such as PRC2 to ensure stable transmission of these chromatin signatures through mitotic division ^41, 42, 43^. However, a role for PRC2 in immune cells as a regulator of metabolic disease has not been previously described. In our study, we found that vitamin D deficiency in utero persistently downregulates *Jarid2* expression and activates the immune cell *Mef2/PGC1α* pathway in donor HSCs, recipient BM, and macrophages despite postnatal VD supplementation, suggesting a stable epigenetic program in immune cells. DNA methylation is one crucial epigenetic mechanism to suppress essential genes during differentiation ^44^. Evidence linking maternal VD status to global DNA methylation in offspring liver, adipose tissue, and leukocytes has been described in multiple animal models and in pregnant women ^25, 45, 46, 47, 48^. In this study, NGS confirmed that vitamin D deficiency promotes methylation of numerous CpG islands in putative enhancers between 5’ upstream to 3’ UTR regions of the mouse *Jarid2* gene. Thus, VD-dependent regulation of *Jarid2* and consequently PRC2 may have the capacity to induce vast changes in the methylome by attenuating the function of major regulator of chromatin structure and function. Future studies using in situ mutagenesis will be critical to confirm which methylated enhancers influence *Jarid2* expression.

PGC1α is a central transcriptional regulator of mitochondrial biogenesis and function and is activated in proinflammatory adipose macrophages in the context of insulin resistance induced by high fat diet ^49^. This study demonstrated that HSCs from vitamin D-deficient fetal donors consistently maintained *Mef2/PGC1α* pathway activation when transplanted into recipients. Moreover, the activated PGC1α network in recipient adipose tissue macrophages promoted a proinflammatory phenotype with increased secretion of miR106b-5p, causing adipose IR. Previous studies indicate that overexpression of PGC1α increases exosome biogenesis and expression of the endosomal sorting complex required for transport (ESCRT), suggesting a role of mitochondrial bioenergetics in macrophage handling of miRNA cargo into exosomes for secretion ^50^. Our previous data demonstrate that selective knockout of the macrophage VDR during embryogenesis promotes miR-106b-5p secretion that activates juxtaglomerular cells to induce hyperreninemic hypertension ^33^. Together, these data identify that the Jarid2/Mef2/PGC1α program induced by vitamin D deficiency in utero enables the communication between innate immune cells, adipocytes, and JG cells to cause cardiometabolic disease.

Recently, microRNAs (miRNAs) have been linked to the regulation of chronic inflammation and IR ^37, 38^. The role of circulating miRNAs as signaling mediators of cell-to-cell communication has been identified both in physiological and pathological conditions ^51^. Multiple miRNAs have been linked directly to regulation of insulin secretion and action on metabolic tissues in diabetic patients and in mouse models of insulin resistance. However, it is notable that miRNAs associated with insulin signaling genes are poorly conserved across species, perhaps implying that those that are conserved may be essential for insulin action and more clinically relevant ^52, 53, 54^. Recent studies show that miR-106b is increased in human circulation in the setting of IR and is highly expressed in the skeletal muscle of both humans and mice ^38, 55, 56^. Moreover, in skeletal muscle cells, overexpression of miR-106b impairs insulin-stimulated glucose uptake and GLUT4 transportation, suggesting an important role of circulating and tissue-specific miR-106b in IR ^57^, However, the cell or tissue type that secretes this miR-106b and the conditions that trigger it have not been previously identified. In this study, we provide evidence via multiple mouse models that macrophages epigenetically programmed in utero by vitamin D deficiency are a source of miR-106b-5p, which induces adipocyte IR. In adipocytes, PIK3CA and PIK3CB are the primary insulin-responsive PIK3 subunits ^58^. Interestingly, mice lacking adipose PIK3CA but not PIK3CB exhibit increased adiposity, glucose intolerance and liver steatosis ^39^. In this study, we provide confirmation via multiple mouse models that VD(−) HSC recipient peritoneal macrophage secrete increased miR-106b-5p, which enters into adipocytes and downregulates PIK3CA and PDK1 expression preventing insulin-stimulated AKT activation and adipose glucose uptake. This finding delineates an unprecedented mechanism by which the miRNA secretome of VD(−)HSC-derived macrophages can regulate adipose insulin signaling.

In summary, our findings provide evidence that an epigenetic immune program in response to in utero vitamin D deficiency is sufficient to cause IR by a miRNA-specific mechanism that enables communication from innate immune cells to adipocytes. This program activates the *Jarid2/Mef2/PGC1α* pathway in immune cells, which persists across both differentiation and transplantation, highlighting the durability of these changes in the offspring regardless of subsequent vitamin D status. Similar alterations in the *Jarid2/Mef2/PGC1α* pathway are replicated in cord blood monocytes from vitamin D deficient mothers. These results identify the need for clinical trials to prove that the widespread screening and treatment of vitamin D deficiency in pregnant women will reduce the long-term risk of cardiometabolic disease in their children and subsequent generations.

## Methods

### Animal Models

Three different mouse transplant models were used: 1) C57BL/6 CD45.1^+^ donors into C57BL/6, CD45.2^+^ recipients; 2) LDL receptor knockout donors expressing GFP^+/−^ (GFP^+/−^LDLR^−/−^) into GFP^−/−^ LDLR^−/−^ recipients; and 3) miR-106b^−/−^ C57BL6 CD45.2^+^ donors into C57BL/6 CD45.1^+^ recipients. To generate fetal liver hematopoietic stem cell (FL-HSC) donors, we obtained C57BL/6 CD45.1^+^ (Jax/Lab 002014), GFP^+/−^ C57BL/6 (Jax/Lab 004353), LDLR^−/−^ (Jax/Lab 002207) and miR-106b^−/−^ CD45.2^+^ (Jax/Lab 008460) mice. We generated GFP^+/−^ LDLR^−/−^ mice by crossing GFP^+/−^ C57BL/6 with LDLR^−/−^ mice. LDLR^−/−^ were initially crossed with mice that constitutively express the green fluorescent protein (GFP^+/−^) as donors to facilitate identification of GFP-positive embryos by UV lamp and assessment of donor bone marrow chimerism in GFP^−/−^ recipients. C57BL/6 CD45.1^+^ were used as donors and C57BL/6 CD45.2^+^ as recipients to facilitate the determination of engraftment without the influence of GFP. There were no differences in insulin resistance phenotype regardless of engraftment verification model. To generate vitamin D-deficient or -sufficient [VD(−) or VD(+)] FL-HSC donors, we transitioned the dam’s diet four weeks prior to pregnancy to either vitamin D-deficient (Harlan TD.87095) or sufficient (Harlan TD.96348) diet ^59^. Females were mated with vitamin D-sufficient males to prevent the effects of vitamin D deficiency on male fertility. At gestational day 13.5, VD(−) or VD(+) FL-HSCs from males and females were harvested for transplantation. A subgroup of pregnant dams was allowed to progress to term. Pups born both vitamin D-deficient and sufficient were weaned to vitamin D-deficient or sufficient diets for 8 weeks. Peritoneal macrophages were obtained for RNA expression. All experiments included male and female animals. Protocols were approved by the Washington University Institutional Animal Care and Use Committee and are compliant with ethical regulations for studies involving laboratory animals.

### Primary FL-HSC transplants and secondary BM transplants

Fetal liver cells at embryonic day 13 include HSCs with a high proliferative capacity, increasing donor engraftment by 10-fold compared to BM stem cell donors ^60^. *For fetal liver transplantation*, vitamin D-deficient or sufficient C57BL6 or LDLR^−/−^ pregnant mice were sacrificed at 13.5 days gestation, with the vaginal plug counted as day 0.5 ^61^. For GFP-model, positive embryos were selected by UV lamp before dissecting out each fetal liver. Fetal livers were then rinsed in sterile saline followed by trypsinization for 15 min at 37 °C. Fetal liver cells were resuspended in cold DMEM with 5% fetal bovine serum (FBS, Gibco), filtered through a 70-µm filter (BD), centrifuged at 125 *g* for 10 min, re-filtered through a 40-µm filter, and centrifuged at 125 *g* for 5 min. The cells were then rinsed in 15 mL of cold phosphate-buffered saline (PBS), pelleted, resuspended in 1mL of PBS, counted using a hemocytometer, and adjusted to 10^5^ cells per µL. Cells were genotyped by PCR for the Sry sex-determining region of chr Y to create mixed male and female pools for donation. Fetal liver cells including HSCs were injected intravenously within 8 hours into 8-week-old vitamin D-sufficient recipient male and female mice following lethal irradiation with 10 Gy from a ^137^Cs gamma irradiator source. *For bone marrow transplantation*, BM cells were isolated from 24-week-post-primarily-transplanted-recipients of VD(−) or VD(+) HSCs by flushing the femurs and tibias with ice-cold PBS ^28^. Total BM was washed, triturated using a 24-gauge needle (Benson Dickson), collected by centrifugation at 300 *g* for 4 min, and diluted with PBS. After lysis of erythrocytes using 0.05% sodium azide, cells were counted. Eight-week-old vitamin D-sufficient C57BL6 (CD45.2^+^) or GFP^−/−^ LDLR^−/−^ male and female recipient mice were lethally irradiated with 10 Gy from a ^137^Cs gamma irradiator source. Within 6 h after irradiation, recipient BM was reconstituted with ∼5 × 10^6^ donor marrow cells via a single injection. Eight weeks after either type of transplantation and reconstitution, recipient engraftment was evaluated by flow cytometry quantification of the percentage of CD45.1 or GFP positivity in peripheral leukocytes of recipients, with only those animals >87% chimeric used in experiments ^28, 33^.

### Metabolic assessment

*Blood Samples*: Fasting serum glucose, cholesterol, triglycerides, and FFA were measured after 6 h of fasting using commercially available kits. For glucose and insulin tolerance tests, transplant recipient mice were evaluated at 8 weeks or 24 weeks post-transplant. Mice were fasted for 6h prior to peritoneal injection with 10% D-glucose (1g/kg) or insulin (0.75 U/kg, Humulin R). For both studies, tail vein blood glucose was assayed using a glucometer at baseline and 30, 60, and 120 min after injection ^62^. Plasma insulin was assessed at 30 min during GTT (by electrochemiluminescence immunoassay). Hyperinsulinemic euglycemic clamps and *in vivo* 2-deoxyglucose uptake (2-DG) assays were performed five days after double lumen catheters were placed. Animals were fasted overnight, and glucose turnover was measured in the basal state and during the clamp at 12-weeks post-transplant in conscious mice as previously described ^28, 62, 63^. Immediately after euthanasia, hind limb muscles and perigonadal fat were harvested, washed with PBS, and placed in liquid nitrogen until pending analysis. Frozen tissue samples were ground, boiled, and centrifuged. Accumulated 2-DG in the supernatant was separated by ion exchange chromatography using a Dowex 1-X8 (100–200 mesh) anion exchange column. Data is expressed as µmol/100 grams of tissue/min [(2-DG x mean blood glucose)/area under the curve]. *<u>For insulin-stimulated adipocyte 2-DG uptake</u>*, differentiated 3T3-L1 adipocytes or primary adipocytes were co-cultured for 72h with peritoneal or stromal vascular macrophages or exposed to described conditioned media. For co-culture, macrophages (0.3 ×10^6^) were placed on inserts in transwell plates with 3T3-L1 adipocytes or primary isolated perigonadal adipocytes in the bottom chamber. After macrophage co-culture, adipocytes were serum-starved, washed, and incubated with or without insulin (10nM) for 30 min, then incubated for 10 min with radioactive 2-DG (PerkinElmer NEC 720A250UC). Cells were washed in cold Krebs-Ringer phosphate HEPES (KRPH), and after lysis, [^14^C] was determined by scintillation counting to measure 2-DG uptake ^64^. Cytochalasin B, an inhibitor of glucose transport (50 µM), was used to correct for non-specific background uptake. Data is presented as a ratio of 2-DG uptake after insulin stimulation to that of non-insulin-stimulated cells. Immunoblots for phospho-AKT (Ser 473) (Cell signaling #4058 dilution 1ug/mL), AKT (Cell signaling #112580 dilution 0.5 ug/mL), and β-Actin (Cell signaling #8457 dilution 0.5 ug/mL) were performed in homogenized perigonadal adipose tissue from HSC recipients with or without insulin stimulation (Humulin R) 1 ug/mL for 5 min. *<u>For skeletal muscle 2-DG uptake</u>*, primarily transplanted mice were fasted for 6h, then paired soleus and extensor digitorum longus muscle (EDL) muscles from anesthetized mice were excised and incubated using a 2-step incubation protocol ^63^. For all incubation steps, vials were continuously gassed with 95% O_2_/5% CO_2_ and shaken in a heated water bath, and one muscle from each mouse was incubated in solution supplemented with 100 μU/ml of insulin (Humulin R) while the contralateral muscle was incubated in solution without insulin (basal) followed by 3 incubation steps as previously described ^63^. After incubation with 2-DG for 15 min, muscles were rapidly blotted on filter paper dampened with incubation medium, trimmed, freeze-clamped, and stored at −80°C. Muscles were homogenized and 2-DG uptake was determined by scintillation counting.

### Isolation of peritoneal and adipose tissue macrophages and primary adipocytes

*Peritoneal macrophages*. Unstimulated macrophages were collected following peritoneal PBS injection and placed in 12-well transwell inserts (Costar polycarbonate filters, 3 µm pore size) for co-culture with adipocytes or in 12-well plates for 6-8h for media collection as previously described ^28, 59^. *Stromal vascular fraction isolation*. Perigonadal fat pads were excised and minced in PBS with 0.5% BSA. Collagenase I (1 mg/ml, Millipore Sigma) was added before incubation with shaking/rotating, and digestion was stopped with pre-warmed KHB (Millipore Sigma). The cell suspension was filtered through a 250-μm filter and spun at 500*g* x 5 min to separate floating adipocytes from the stromal vascular fraction (SVF) pellet. The SVF pellet was resuspended in FACS buffer and incubated with anti-CD14 magnetic microbeads (Miltenyi Biotec) and F4/80 to isolate adipose tissue macrophages (ATM), and ATM macrophages were placed in 12-well plates with transwell inserts for adipocyte culture or in collagen-coated plates with DMEM plus 10% exosome-depleted FBS for 6h for media miRNA expression and cytokine analysis ^65^. *Primary adipocytes*. Perigonadal fat pads were initially treated with collagenase as described above, but then floating adipocytes were spun at 100 *g* x 1 min. Buffer and SVF pellet underneath the floating adipocytes were removed, and cells were washed with prewarmed KRPH. Adipocytes were transferred into collagen-I-coated 12-well plates (0.5 ×10^6^ cells/plate) and incubated at 37°C for 60 min before starting the 2-DG protocol.

### 3T3-L1 adipocyte differentiation

Murine 3T3-L1 pre-adipocytes (American Type Culture Collection) were grown, maintained, and induced to differentiate using standard protocol (<u>7</u>). Fully differentiated adipocytes (12 days post differentiation induction) were maintained in DMEM supplemented with 10% FBS (Millipore Sigma) until two days before experimentation when cells were fed with 10% calf serum (Millipore Sigma). Prior to experiments, media was changed to serum-starved DMEM (low glucose) for 2-3 hr.

### Macrophage and adipocyte co-cultures

Transwell chambers were utilized (Costar polycarbonate filters, 3 µm pore size) as previously described ^28^ for co-cultures. Membranes and 12-well plates were coated with fibronectin (Life Technologies) overnight at 4 degrees. Peritoneal or SVF macrophages were cultured in DMEM plus 10% exosome-depleted FBS for 6 hrs, and media was tested for miRNA expression and cytokine levels prior to co-culture. Differentiated 3T3-L1 adipocytes were grown to 80-90% confluence in 12-well plates, maximum 0.5 × 10^6^ cells/well. Macrophages (0.3 × 10^5^ cells/well) were added to the transwell upper chamber, with adipocytes in the lower chamber. Cells were co-cultured for 72 h in DMEM/F12 media with 10% exosome-depleted FBS with 2-DG quantification as described above. Some 3T3-L1 cells were cultured only with macrophage conditioned media, but additionally incubated with 0.2 µg/ml TNFα-, IL-1β-, or IL-6-neutralizing antibodies (R&D Biosystems MAB4101) for 72 h, with 2-DG quantification as described above ^63^.

### Flow cytometry

Monocyte and SVF macrophages cell surface marker analysis was performed using a FACStar Plus as previously described ^66, 67^. After isolation, including CD11b selection with microbeads, cells were resuspended in flow cytometry buffer, and >10^5^ cells were analyzed for each sample. Monocytes were incubated with 0.2 mg/mL of anti-mouse APC-CD45.1 (BioLegend, #110714) or anti-mouse PE-CD45.2 (BioLegend #109808) for 15 min on ice, then washed before flow cytometry with utilization of APC mouse IgG2a/K (BioLegend #400219) and PE mouse IgG2a/K (BioLegend #400211) isotype controls. EGFP positive cells were excited at 488 nm and measured by flow cytometry at 530 nm. Cell aggregates, dead, and cellular debris were excluded based on FSC/SCC. CD11b+-SVF were incubated with were with 0.2 mg/mL of anti-mouse PE-F4/80 (BioLegend, #12-4801-82) with utilization of PE rat IgG2a K isotype Control (BioLegend#12-4321-80) to determine the percentage of ATM cells. Batch analysis by Flow Jo version 9.6.2 was used for gating consistency and selection of positive populations. Unstained samples and blocking with FC were used to decrease autofluorescence and non-specific background. Flow cytometry data is presented as the percentage of fluorophore-or GFP-positive live cells.

### F4/80 immunofluorescence staining

For tissue sections, ketamine/xylazine-anesthetized mice were perfused for 10 min with 4% paraformaldehyde before perigonadal fat pads were collected and immersed in the same fixative for 12–15 h at 4°C. After rinsing and PBS wash, tissue was dehydrated with gradual steps of EtOH and paraffin-embedded. Adipose tissue was cut into 3–4 μm sections, then slides were deparaffinized, rehydrated, and blocked for endogenous peroxidase activity (1% H_2_O_2_ in TBST). Slides were stained with F4/80-specific antibodies (1:300 Abcam ab6640) following the manufacturer’s recommendations. F4/80-positive cells were counted in 15 different fields with a 20x objective.

### Microarray and bioinformatics analysis

Purified DNA-free RNA from primary recipients BM cells, was quantified and quality was assessed using a Nanodrop ND-100 spectrophotometer and hybridized to Affymetrix GeneChip miRNA 4.0 microarrays by Washington University’s Genome Technology Access Center. Postprocessing of array signal data was performed with Partek Genomics Suite 6.6 (Partek, St Louis, MO). Upregulated genes (Ratio >1.19 and nominal p<0.05) and downregulated genes (Ratio <0.94 and nominal p<0.05) in BM of VD-FL-HSC recipients were used as input for enrichment pathway analysis using Enrichr, Embryonic Stem Cells Atlas of Pluripotency Evidence (ESCAPE), gene Ontology (GO), Kyoto Encyclopedia of Genes and Genomes (KEGG), and Wiki-pathway databases. RNA sequences from bone marrow arrays data (GSE158763) have been deposited in the NCBI GEO repository.

### Methylation analysis

Bone marrow DNA was isolated, and next generation sequencing methylation analyses were performed by EpigenDx, Inc. (Hopkinton, Massachusetts, United States). 500 ng of extracted DNA samples were bisulfite modified using the EZ-96 DNA Methylation-Direct KitTM (ZymoResearch; Irvine, CA; cat# D5023) per the manufacturer’s protocol. All bisulfite modified DNA samples were amplified using separate multiplex or simplex PCRs. PCR products from the same sample were pooled, and libraries were prepared using a custom Library Preparation method created by EpigenDx. Library molecules were purified using Agencourt AMPure XP beads (Beckman Coulter; Brea, CA; cat# A63882). Barcoded samples were then pooled in an equimolar fashion before template preparation and enrichment were performed on the Ion ChefTM system using Ion 520TM & Ion 530TM ExT Chef reagents (Thermo Fisher; Waltham, MA; cat# A30670). Following this, enriched, template-positive library molecules were sequenced on the Ion S5TM sequencer using an Ion 530TM sequencing chip (cat# A27764). FASTQ files from the Ion Torrent S5 server were aligned to the local reference database using open-source Bismark Bisulfite Read Mapper with the Bowtie2 alignment algorithm. Methylation levels were calculated in Bismark by dividing the number of methylated reads by the total number of reads. The % methylation increase was calculated as CpG methylation of recipients of HSC VD(−) -CpG methylation of HSC VD(+) recipients/ CpG methylation of HSC VD(+) recipients.

### Exosome isolation

Media was centrifuged at 2000 *g* for 30 minutes. Supernatant was transferred to a new tube and centrifuged at 100,000 *g* for 18 h. Density-gradient-based isolation was then performed using the Total Exosome Isolation kit (Invitrogen) ^68^. The purity of exosome vesicles was confirmed by Western blot showing the presence of EV marker protein expression as previously described ^33, 68^.

### MicroRNA and mRNA expression via RT-qPCR

miRNA purification and isolation from 3T3-L1 cells was performed using the *mir*Vana miRNA kit (Invitrogen Ambion) and anti-miR (AM10067). RTq-PCR was performed using the TaqMan reagent kit, with relative expression of miRNA calculated by the comparative threshold cycle method relative to miR-39 from *C. elegans*. For media assays, we spiked in miR-39 as an exogenous housekeeping miRNA control prior to extraction (Qiagen #219610). TaqMan primers were obtained from Life Technologies for miRNAs 106b 5p (#000442), 106b-3p (#002380), let7g-5p (#002282), 142-3p (#000464), 330-3p(#001062), 19b-5p (#002425), 125a-5p (#002198), 331-3p (#000545), U6 (#4427975) and 39 (#467942). mRNA RTq-PCR was performed with the GeneAmp 7000 Sequence Detection System using the SYBR® Green reagent kit (Applied Biosystems)^33^. We used the following mouse oligonucleotides; for *Jarid2* forward 5′-GCGGTAAATGGGCTTCTTGG-3’; *Jarid2* reverse 5’-TGCTAGTAGAGGACACTTGGGA-3’; *Ppargc1a* forward 5′-AGCCTCTTTGCCCAGATCTT-3’; *Ppargc1a* reverse 5′-GGCAATCCGTCTTCATCCAC-3’, *Mef2b* forward 5′-GACCGTGTGCTGCTGAAGTA-3; *Mef2b* reverse 5′-AGCGT CTCGAGGATGTCAGT-3’; *Nrf1* Forward 5′-GTACAAGAGCATGATCCTGGA-3’; *Nrf1-*reverse 5′-GCTCTTCTGTGCGGACATC-3’; *Atp5g* forward 5’-AGTTGGTGTGGCTGGATCA-3’; *Atp5g* reverse 5’-GCTGCTTGAGAGATGGGTTC-3’; *Pparb* forward 5′-TGGAGCTCGATGACAGTGAC-3’; *Pparb* reverse 5′-GGTTGACCTGCAGATGGAAT-3’;

*Fabp5* forward 5’-AGAGCACAGTGAAGACGAC-3’; *Fabp5* reverse 5’-CATGACACACTCCACGATCA-3’; *Mrpl32* forward, 5’-AAGCGAAACTGGCGGAAAC-3’; *Mrpl32* reverse, 5’-GATCTGGCCCTTGAACCTTCT-3’; *PDPK1* forward, 5’-CCACTGAGGAAGATCGACAGAC-3’; *PDPK1* reverse, 5’-AGAGGCGTGATATGGGCAATCC-3’; *PIK3CB* forward, 5’-CAGTTTGGTGTCATCCTGGAAGC-3’; *PIK3CB* reverse, 5’-TCTGCTCAGCTTCACCGCATTC-3’; *PIK3CA* forward, 5’-CACCTGAACAGACAAGTAGAGGC-3’; *PIK3CA* reverse, 5’-GCAAAGCATCCATGAAGTCTGGC-3’.All assays were done in triplicate, with data expressed as relative expression of mRNA normalized to mouse ribosomal protein *Mrpl32*.

### Plasmids and small interfering RNA transfection

3T3-L1 cells were transfected with mirVana miR106b (LifeTechnology MC10067), let 7g-5p (LifeTechnology MC11758), 330-3p (LifeTechnology MC10732), 19b-5p (LifeTechnology MC13042), 142-3p (LifeTechnology M C10398),125a-5p (LifeTechnology MC12561), 331-3p(LifeTechnology MC10881) mimics or miR106b antagomir (AM10067). 72 h after transfection, 3T3-L1 cells were evaluated for 2-DG uptake and mRNA expression in lysates by RT-qPCR. 3T3-L1 cells were also transfected with pre-miR-106b siRNA sense oligonucleotides 5’-CCU AAU GAC CCU CAA GCC GUU-3 and antisense 5’-CGG CUU GAG GGU CAU UAG GUU-3’ then exposed to VD(+) HSC or VD(−) HSC recipient peritoneal macrophage media for 72 h before assessment of pre-miR-106-5p or mature miR-106b-5p. Peritoneal macrophages from VD(+) HSC or VD(−) HSC recipients were transduced with lentivirus containing sense either Jarid2-siRNA (LifeTechnology 4390771), *Ppargc1a*-siRNA (LifeTechnology, AM16708), or control-siRNA (LifeTechnology AM461) for 48 hours and mRNA expression was determined as previously described ^69^.

### Western Blot Analysis

3T3-L1 adipocytes were homogenized in RIPA lysis buffer containing protease and phosphatase inhibitors. Lysates were clarified, centrifuged, and resolved by SDS-PAGE. Samples were transferred to PVDF membranes that were subsequently probed with the following antibodies for protein and phosphoprotein detection: anti-PIK3CA (ab40776, Abcam), anti-PDPK1 (ab52893, Abcam,) ant-AKT (9272S, Cell Signaling Technology), anti-pAKT (9271L, Cell Signaling Technology) and β-actin (3700S, Cell Signaling Technology). Protein levels were quantified using Image Studio Lite Ver 5.2 (LI-COR) and normalized to β-actin protein levels.

### Human Samples

Specimens & data were obtained in a de-identified manner from the Women and Infant Specimen Consortium (WIHSC) IRB#201013004, consisting of 30 healthy pregnant women prior to delivery and their full-term infants. Women with singleton pregnancies resulting in full-term vaginal or C-section delivery were included, and intrauterine growth retardation, small/large for gestational age, diabetes (gestational, type 1, and type 2 DM), preeclampsia, chorioamnionitis, acute infection (fever or active herpes), and moderate or severe alcohol or drug abuse during pregnancy were excluded. Venous cord blood plasma was collected for 25-hydroxyvitamin D levels (LC-MS/MS) ^66, 67^ and quantification of miRNA expression from plasma exosomes ^68, 70^. Cord blood monocytes were isolated as previously described ^69^. Monocytes were stabilized for 2 hours in 100% serum from the original patient to mimic in vivo conditions for experiments^66, 67^. Isolated monocytes were co-cultured with 3T3-L1 adipocytes in transwell chambers for insulin-stimulated 2-DG studies. Monocyte mRNA was also assessed by qPCR^69^.

### Statistical Analysis

Experiments were carried out in duplicate or triplicate, with “n” referring to the number of distinct samples. Gaussian distribution was verified by Kolmogorov-Smirnov distance. Parametric data are expressed as mean ± SEM and analyzed by two-sided t-tests, paired or unpaired as appropriate, or by one-way ANOVA and Tukey’s post-test for more than two groups. Statistical analysis was carried out using GraphPad Prism version 8.4.3.

## Data and materials availability

RNA sequences from bone marrow arrays data (GSE158763) have been deposited in the NCBI GEO repository

## Computer Code Availability

There is no custom computer code or algorithm used to generate results in this manuscript.

## Acknowledgments

This work was supported by NIH UL1TR002345 (CBM, AER, KPM, JO), NIH P30DK020579 (CBM), VA Merit Award 1BX003648-01 (CBM, JO, AER, AD, MS, KTB, RAB, RDH) Child Discovery Institute CH-II-2012-209 (CBM, AER, KPM, TW, XX, DL, JO, JS) The contents of this article are solely the responsibility of the authors and do not necessarily represent the official view of NIH.

## Author contributions

J.O., A.E.R., K.P.M., I.D., R.M.Z., R.D.H., R.A.B., K.T.B., A.D., J.S., T.W.,C.M., M.B., M.M., A.C.,M.S.,C.B.-M. planned the experiments. J.O., I.D., J.S., M.S. C.M., performed animals transplants and metabolic assessment. R.D.H and R.A.B performed and analyzed the gene array experiments. T.W., C.M., J.S., X.X., D.L., performed and analyzed the methylation experiments. A. E.R., K.T.B., A.D., K.P.M., R.M.Z., performed and analyzed human monocyte experiments. C.BM. wrote and revised the paper. All authors have given approval to the final version of the paper.

## Additional Information

Supplementary Figures 1-6

Supplementary Table 1-4

