## Supplemental Fig and tables for "Embryonic Vitamin D Deficiency Programs Hematopoietic Stem Cells to Induce Type 2 Diabetes"

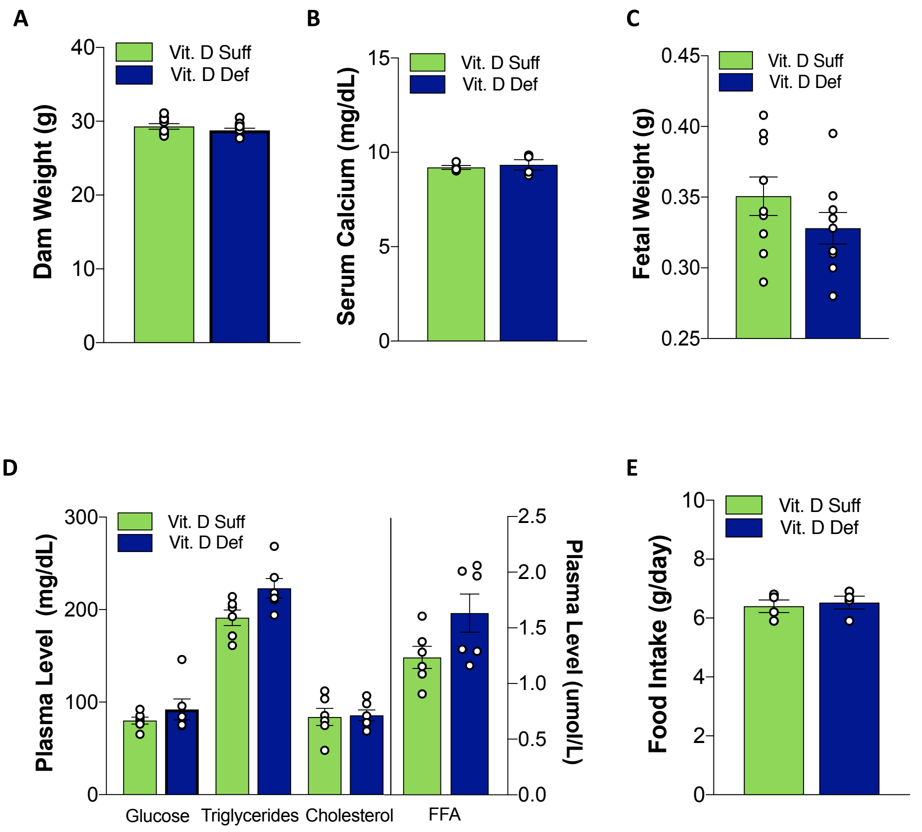


**Fig. S1. Maternal vitamin D deficiency during pregnancy does not alter metabolic parameters in dams. (A)** Dam weight (n=9 vitamin D sufficient, n=10 vitamin D deficient), **(B)** dam serum calcium (n=4 per group), **(C)** fetal weight (n=7 per group), **(D)** dam plasma fasting glucose and lipids (n=6 per group), and **(E)** dam food intake (n=4 per group)in vitamin D-sufficient or -deficient mice pregnant at 13 days and their fetuses. Data presented as mean ± SEM.

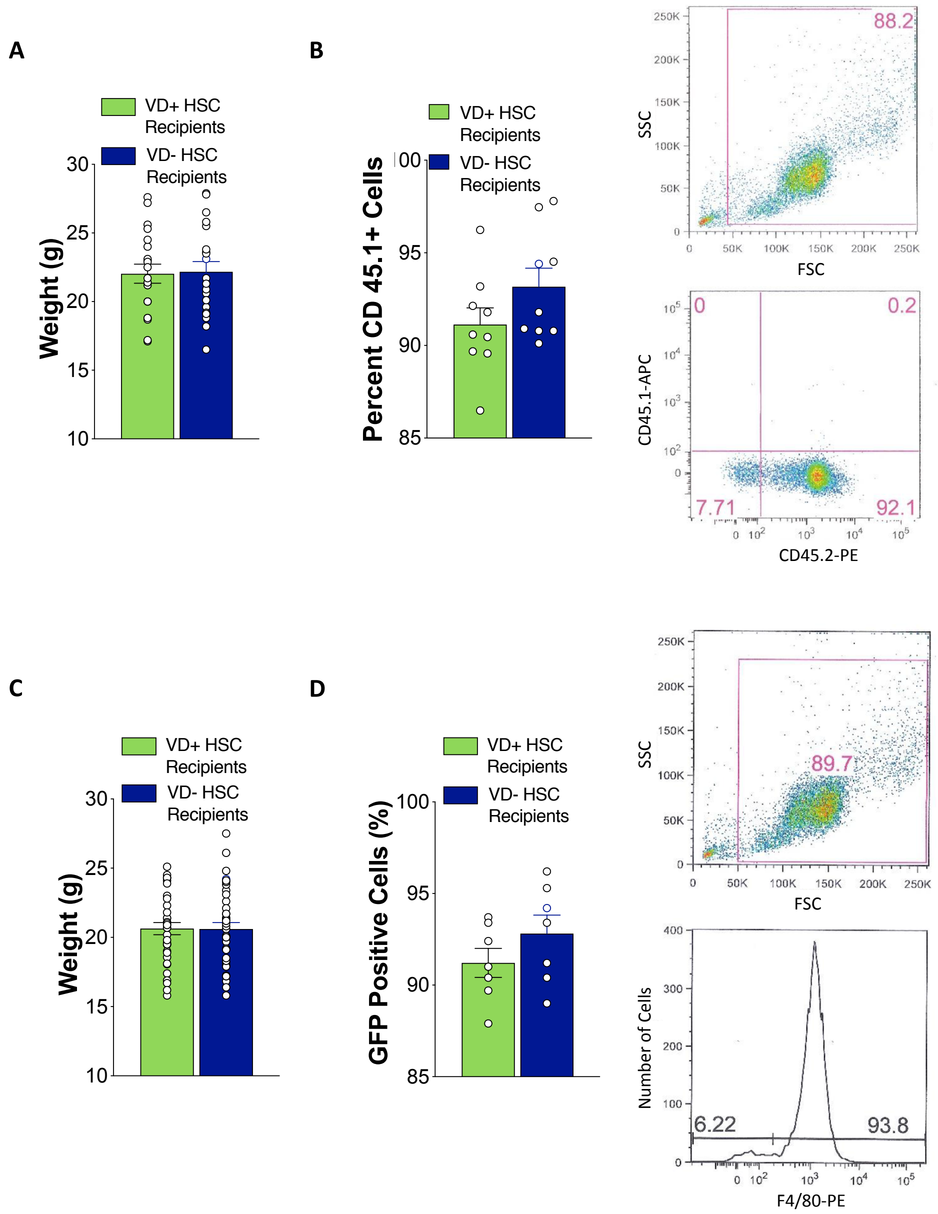


**Fig. S2. Vitamin D status of HSCs does not alter recipient weight or engraftment.** HSCs isolated from embryos exposed to in utero vitamin D sufficiency or deficiency were transplanted into vitamin D-sufficient recipient mice in C57BL6 and LDLR$^{-/-}$ backgrounds. **(A)** Weight (n=20 per group) and **(B)** percentage of circulating 45.1-positive cells (n=9 per group) was measured 8 weeks after transplantation into C57BL6 (CD45.2$^{+}$) recipient mice. **(C)** Weight (n=35 VD+, n=/38 VD-) and **(D)** percentage of circulating GFP-positive cells (n=7 per group) was measured 8 weeks after transplantation in LDLR$^{-/-}$ GFP$^{-/-}$ recipient mice. Data presented as mean ± SEM.

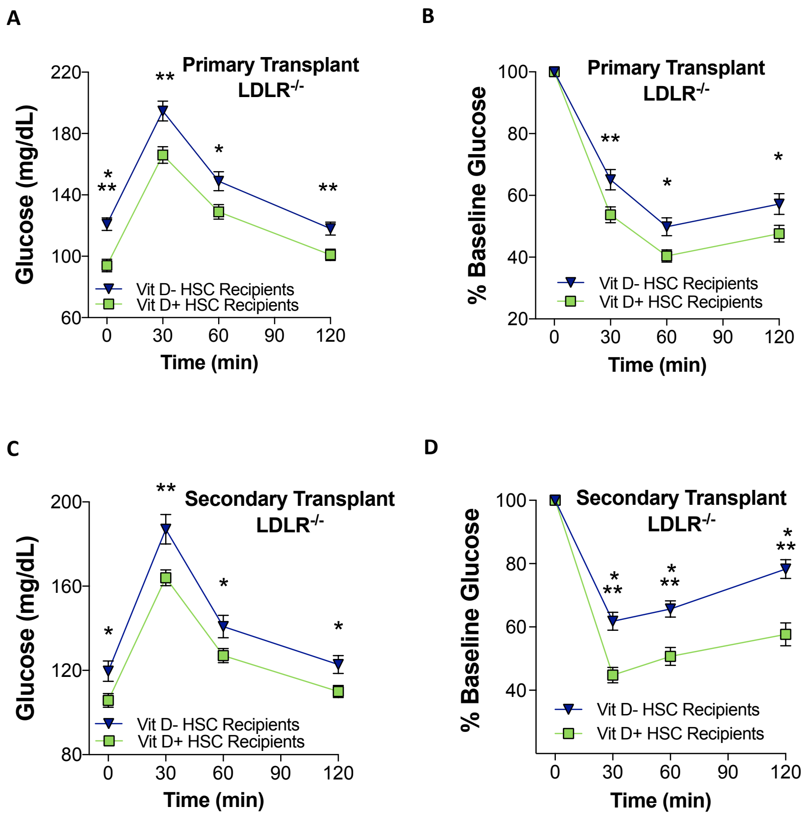


**Fig. S3. In utero vitamin D deficiency reprograms HSCs to transfer insulin resistance in the LDLR$^{-/-}$ background.** Vitamin D sufficient LDLR$^{-/-}$ GFP$^{-/-}$ recipient mice were transplanted with VD(-) or VD(+) FL-HSCs from LDLR$^{-/-}$ GFP$^{+/-}$ (primary transplant). Primary recipient BM was then used as a transplant donor for vitamin D sufficient mice (n=38-42/group) (secondary transplant). Glucose and insulin tolerance tests were performed at **(A, B)** 8 weeks and **(C-D)** 8 weeks post-secondary-transplant. Data presented as mean ± SEM. *p<0.05, **p<0.01, ***p<0.001 vs. primary or secondary recipients of VD(+) HSCs by two-tailed unpaired t test.

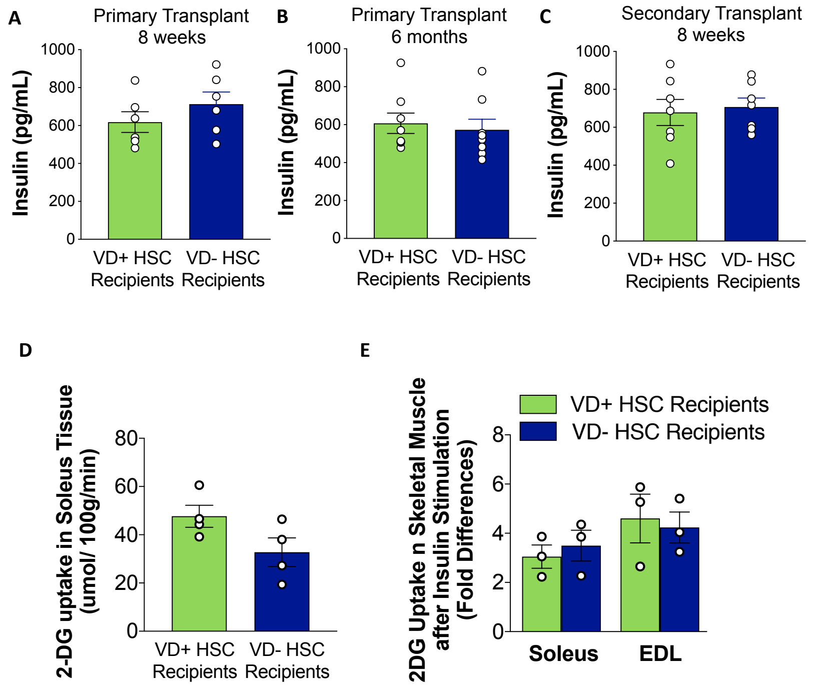


**Fig. S4. Recipient insulin levels after glucose challenge and skeletal muscle insulin resistance are not affected by vitamin D status of HSCs.** Vitamin D sufficient CD45.2$^+$ C57BL6 mice were transplanted with VD(-) or VD(+) FL-HSCs from C57BL6 mice CD45.1$^+$ (primary transplant). Primary recipient BM was then used as a transplant donor for vitamin D sufficient mice (secondary transplant). Plasma insulin levels were measured 30 min after glucose challenge **(A)** 8 weeks after transplantation (n= 6 per group), **(B)** 6 months after transplantation (n=8 per group), and **(C)** 8 weeks after secondary transplantation (n=7 per group). Skeletal muscle insulin sensitivity was measured in primary recipients 8 weeks post-transplant by **(D)** 2-deoxyglucose (2-DG) uptake in soleus muscle during hyperinsulinemic euglycemic clamps (n=4 per group) and **(E)** insulin-stimulated 2-DG uptake by ex vivo soleus and extensor digitorum longus muscle (EDL) (n=3 per group). Data presented as mean ± SEM. *p<0.05, **p<0.01, ***p<0.001.

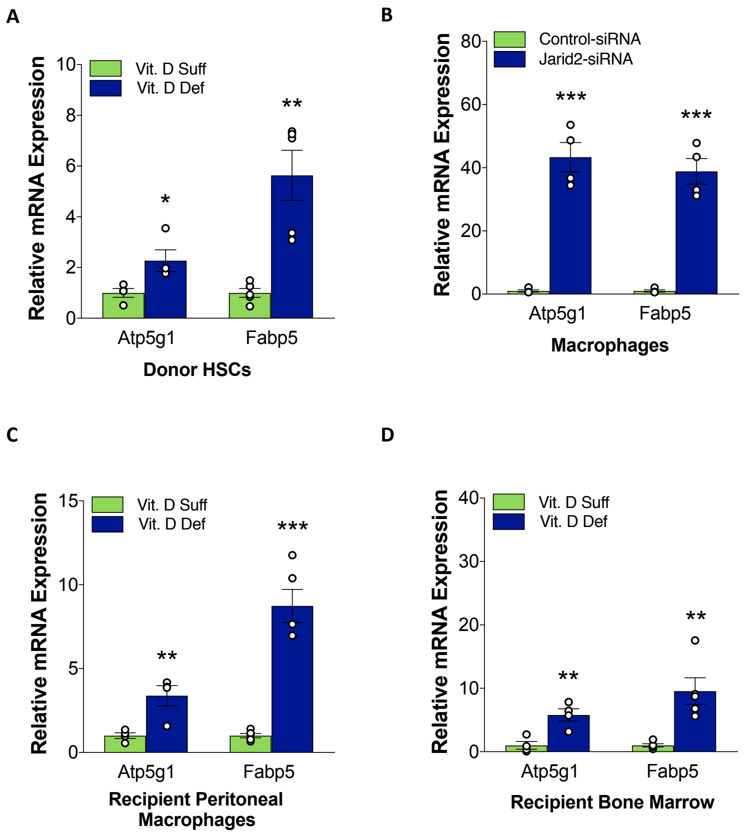


**Fig. S5. In utero vitamin D deficiency activates the target genes of the Jarid2/Mef2/PGC1$\alpha$ network in immune cells.** Quantitative RT-PCR (n=4/group) in **(A)** donor FL-HSCs, **(B)** Quantitative RT-PCR (n=4/group) in peritoneal macrophages transfected with Jarid2-siRNA vs. control-siRNA to analyze target genes of the Jarid2 and PGC1$\alpha$ network. **(C)** BM, and **(D)** peritoneal macrophages from FL-HSC transplant recipients to analyze target genes of the Jarid2 and PGC1$\alpha$ network. Data presented as mean ± SEM. *$p<0.05$, **$p<0.01$, ***$p<0.005$ vs. VD (+) HSC donors or recipients or control-siRNA by two-tailed unpaired t test.

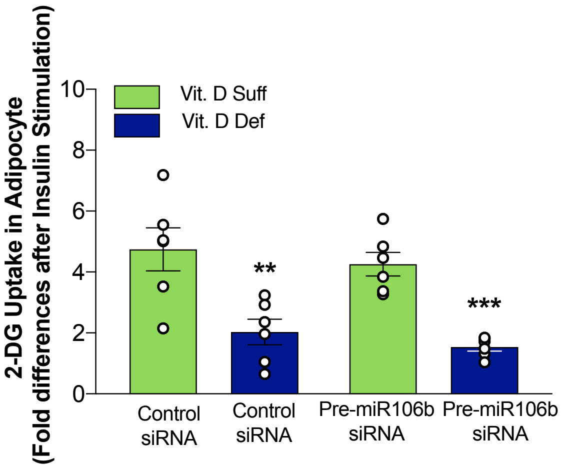


**Fig. S6. Endogenous miR-106b-5p production by the adipocyte is not responsible for the impaired 2-DG uptake.** Insulin-stimulated 2-DG uptake in 3T3-L1 adipocytes transfected with pre- miR-106b-siRNA or control-siRNA and exposed to conditioned media from peritoneal macrophages from VD(+) and VD(-) FL-HSCs recipients (n=6 per group). Data presented as mean ± SEM. **p<0.01, ***p<0.005.

### Enrichr Pathway Analysis

#### Top Up Regulated Pathways

| Term (ESCAPE DATABASE) | Overlap # Genes | P-value | Adjusted P-value |
|---|---|---|---|
| JARID2-20075857 DOWN | 42/1107 | 2.93E-06 | 9.22E-04 |
| CHiP MYC-18555785 | 40/1200 | 9.59E-05 | 0.015107269 |
| CHiP MYC-19079543 | 44/1458 | 3.87E-04 | 0.040608705 |
| mESC H3K36me3 18692474 | 66/2469 | 4.14E-04 | 0.032576428 |
| CHiP CHD1-19587682 | 27/843 | 0.002365341 | 0.149016486 |

| Term (WikiPathways) | Overlap # Genes | P-value | Adjusted P-value |
|---|---|---|---|
| Macrophage markers WP2271 | 3/10 | 7.97E-04 | 0.140340284 |
| Purine metabolism WP2185 | 8/171 | 0.019228259 | 1 |
| Nucleotide GPCRs WP207 | 2/12 | 0.021998268 | 1 |
| Keap1-Nrf2 WP1245 | 2/14 | 0.0295646 | 1 |
| Adipogenesis genes WP447 | 6/134 | 0.047663558 | 1 |

#### Top Down Regulated Pathways

| Term (GO Biological Process) | Overlap # Genes | P-value | Adjusted P-value |
|---|---|---|---|
| Viral myocarditis | 11/87 | 1.28E-04 | 0.03889199 |
| Autoimmune thyroid disease | 9/78 | 0.00101947 | 0.15445045 |
| Allograft rejection | 8/63 | 0.00102296 | 0.10331917 |
| Graft-versus-host disease | 7/64 | 0.00490449 | 0.37151542 |
| Type I diabetes mellitus | 7/69 | 0.00740514 | 0.44875171 |
| Epstein-Barr virus infection | 15/229 | 0.00892602 | 0.45076386 |
| Viral carcinogenesis | 15/229 | 0.00892602 | 0.38636902 |

| Term (KEGG Pathways) | Overlap # Genes | P-value | Adjusted P-value |
|---|---|---|---|
| Positive regulation of stress-activated MAPK cascade (GO:0032874) | 9/80 | 0.00122346 | 1 |
| Regulation of membrane depolarization (GO:0003254) | 4/16 | 0.00153194 | 1 |
| Positive regulation of early endosome to late endosome transport (GO:2000643) | 3/8 | 0.00174629 | 1 |
| Regulation of JNK cascade (GO:0046328) | 9/90 | 0.0027913 | 1 |
| Positive regulation of JNK cascade (GO:0046330) | 8/76 | 0.00343504 | 1 |

**Table S1. EnrichR pathway analysis.**

### Methylation Assay Coordinates

| Assay ID | Assay Location | From ATG | From TSS | GRCm38 (+) | # of CpG |
|---|---|---|---|---|---|
| ASY3159 | 5'-Upstream | -2453 to -2351 | -1852 to -1750 | Chr13:44729419-44729521 | 6 |
| ASY3160 | 5'-Upstream | -2211 to -2124 | -1610 to -1523 | Chr13:44729661-44729748 | 9 |
| ASY3162 | 5'-Upstream | -1157 to -1122 | -556 to -521 | Chr13:44730715-44730750 | 4 |
| ASY3163 | 5'-Upstream | -704 to -649 | -103 to -48 | Chr13:44731168-44731223 | 6 |
| ASY3164 | Exon 1, Intron 1 | 45 to 89 | 646 to 690 | Chr13:44731916-44731960 | 2 |
| ASY3166 | Intron 1 | 2149 to 2235 | 2750 to 2836 | Chr13:44734020-44734106 | 9 |
| ASY3167 | Intron 1 | 35922 to 35937 | 36523 to 36538 | Chr13:44767793-44767808 | 2 |
| ASY3168 | Intron 1 | 39152 to 39183 | 39753 to 39784 | Chr13:44771023-44771054 | 3 |
| ASY1677 | Intron 1 | 42317 to 42469 | 42918 to 43070 | Chr13:44774188 -44774340 | 4 |
| ASY1678 | Intron 1 | 42675 to 42679 | 43276 to 43280 | Chr13:44774546-44774550 | 2 |
| ASY3169 | Intron 1 | 48158 | 48759 | Chr13:44780029 | 1 |
| ASY3170 | Intron 2 | 108192 to 108219 | 108793 to 108820 | Chr13:44840063-44840090 | 2 |
| ASY3173 | Intron 2 | 110157 to 110181 | 110758 to 110782 | Chr13:44842028-44842052 | 2 |
| ASY3175 | Intron 2 | 112064 to 112117 | 112665 to 112718 | Chr13:44843935-44843988 | 3 |
| ASY3176 | Intron 3 | 119607 to 119651 | 120208 to 120252 | Chr13:44851478-44851522 | 2 |
| ASY3178 | Intron 3 | 125037 to 125120 | 125638 to 125721 | Chr13:44856908-44856991 | 3 |
| ASY3179 | Intron 4 | 148554 | 149155 | Chr13:44880425 | 1 |
| ASY3180 | Intron 5 | 158766 | 159367 | Chr13:44890637 | 1 |
| ASY3181 | Intron 5 | 159670 to 159719 | 160271 to 160320 | Chr13:44891541-44891590 | 2 |
| ASY3184 | Intron 7 | 172605 | 173206 | Chr13:44904476 | 1 |
| ASY3185 | Intron 8 | 174879 to 174908 | 175480 to 175509 | Chr13:44906750-44906779 | 2 |
| ASY3186 | Intron 9 | 176115 to 176136 | 176716 to 176737 | Chr13:44907986-44908007 | 2 |
| ASY3188 | Exon 10 | 178525 to 178563 | 179126 to 179164 | Chr13:44910396-44910434 | 3 |
| ASY3190 | 3'-UTR | 188691 to 188728 | 189292 to 189329 | Chr13:44920562-44920599 | 3 |

**Table S2. Next generation sequencing methylation assay coordinates.**

### List of down regulated microRNAs in Recipients Bone Marrow Cells

| Gene Symbol | Mean expression recipients of VD- FL-HSC | Mean expression recipients of VD+ FL-HSC | Ratio Recipient VD- FL-HSC /VD+ FL-HSC | P Value |
|---|---|---|---|---|
| Mir19b-1 | 8.35 | 14.53 | 0.57 | 2.64E-03 |
| Mir106b | 9.83 | 14.60 | 0.67 | 1.12E-02 |
| Mir142 | 80.10 | 111.27 | 0.72 | 2.85E-02 |
| Mir331 | 9.85 | 13.76 | 0.72 | 2.21E-02 |
| Mirlet7g | 13.20 | 18.00 | 0.73 | 1.76E-02 |
| Mir340 | 23.68 | 32.16 | 0.74 | 1.29E-02 |
| Mir330 | 32.60 | 41.23 | 0.79 | 4.36E-03 |
| Mir376b | 3.49 | 4.33 | 0.80 | 2.72E-02 |
| Mir666 | 53.52 | 58.36 | 0.92 | 3.02E-02 |
| Mir679 | 25.32 | 28.72 | 0.88 | 3.32E-02 |
| Mir467b | 3.49 | 3.85 | 0.91 | 3.66E-02 |
| Mir125a | 24.14 | 27.91 | 0.86 | 4.57E-02 |
| Mir92b | 73.49 | 81.69 | 0.90 | 1.08E-02 |

**Table S3. Down-regulate miRNAs from HSC transplant bone marrow.**

### Demographic Characteristics of Population at Delivery

| Maternal Characteristics | (N=30) |
|---|---|
| Age in years (Mean±SE) | 28.4 ± 1.1 |
| African American (%) | 50 |
| Caucasian (%) | 47 |
| Asian (%) | 3 |
| Pre-pregnancy BMI (Mean±SE) | 27.7 ± 1.32 |
| Pre-pregnancy BMI (Mean±SE) | 32.6 ± 1 |
| Vaginal Delivery (%) | 53 |
| C-Section Delivery (%) | 47 |
| Prenatal Vitamin Supplementation (%) | 96% |
| **Infant Characteristics** | |
| Gender Females (%) | 57 |
| Birth Weight (g) (Mean±SE) | 3422 ± 81 |
| Birth Height (cms) (Mean±SE) | 50.7 ± 0.36 |

**Table S4. Demographic characteristic of population at delivery.**
